# A molecular convergence in the triad of Parkinson’s disease, depressive disorder and gut health is revealed by the inflammation-miRNA axis

**DOI:** 10.1101/2025.10.05.680533

**Authors:** Lluis Miquel-Rio, Judith Jericó-Escolar, Claudia Yanes-Castilla, Unai Sarriés-Serrano, Verónica Paz, Luis F Callado, J Javier Meana, Analia Bortolozzi

**Author notes:** Contributed equally to the manuscript as the first authors.

## Abstract

**Background:** Parkinson’s disease (PD) is a multisystem disorder frequently comorbid with non-motor symptoms like depressive disorder (DD) and gastrointestinal (GI) dysfunction. Chronic neuroinflammation and disruption of the gut-brain axis are implicated as shared pathological drivers, but the precise molecular mechanisms connecting these conditions remain elusive. We hypothesized that a common microRNA (miRNA)-mediated inflammatory profile underlies this clinical triad, representing a point of pathological convergence.

**Methods:** We analyzed the expression of a panel of inflammatory bowel disease (IBD)-associated miRNAs, key inflammatory markers, and glial response in postmortem brain tissue (dorsolateral prefrontal cortex and caudate nucleus) from patients with PD, DD, and matched healthy controls. To investigate causality and gut-brain axis involvement, two mouse models were used: (i) PD-associated α-synucleinopathy was induced in dorsal raphe serotonin (5-HT) neurons; and (ii) DD-like based on corticosterone (CORT)-induced stress. Mice were assessed for depressive-like behaviors and GI dysmotility, and their brain (medial prefrontal cortex and caudate-putamen) and ileum tissues were analyzed for the same molecular markers.

**Results:** We identified a conserved miRNA pattern in the brains of both PD and DD patients, characterized by the significant downregulation of miR-199a-5p and miR-219a-5p and the upregulation of miR-200a-3p. This dysregulation was strongly associated with a pro-inflammatory state, as evidenced by increased expression of TNFα, IFN-γ, and NFκB1, as well as changes in the glial response. Mice with α-synucleinopathy in the 5-HT system exhibited a depression-like phenotype and reduced intestinal motility, accompanied by increased Iba1 and GFAP signal. Comparable effects were observed in mice subjected to CORT-induced stress. Notably, the same pattern of miRNAs and inflammatory cytokines observed in the human brain was replicated in the brain and ileum of DD-PD-like mice, providing direct evidence of a parallel pathological process spanning the gut-brain axis.

**Conclusion:** This study identifies a specific miRNA-inflammatory axis as a common molecular mechanism connecting the pathophysiology of PD, DD, and gut dysfunction. This pattern represents a critical point of convergence that drives a shared, bidirectional inflammatory cascade along the gut-brain axis. Targeting this miRNA triad could provide a new therapeutic approach for addressing the motor, psychiatric, and GI symptoms of these interconnected disorders simultaneously.

## Introduction

Parkinson’s disease (PD) is a multisystem disorder, and although clinically characterized by motor symptoms such as bradykinesia, rigidity, and resting tremor, a wide range of disabling non-motor symptoms also occur throughout the course of the disease [1, 2]. Notably, neuropsychiatric symptoms, including anxiety and depression, affect approximately 50% of individuals with PD, and over 80% report gastrointestinal (GI) disorders. These debilitating comorbidities are prevalent across all stages of the disease, from the untreated premotor and early phases to advanced stages [3–8]. The fact that these non-motor symptoms often emerge before the onset of characteristic neurological signs highlights a critical window for early intervention. Despite this, current treatments primarily focus on motor symptom control and do not address disease progression, underscoring the urgent need for disease-modifying therapies.

Recent findings have highlighted the pivotal role of the gut-brain axis in PD pathogenesis, leading to the conceptualization of distinct disease subtypes: "body-first" and “brain-first”, differentiated by the initial site of α-synucleinopathy [9–11]. In the body-first subtype, studies in post-mortem human tissue samples and animal models suggest an initial pathological accumulation of α-synuclein (α-Syn) protein within the enteric nervous system, with subsequent propagation to the dorsal motor nucleus of the vagus (DMV), brainstem, and the rest of the brain [12]. This ascending pathology affects serotonin (5-HT) neurons in the caudal raphe nuclei early in the disease course, preceding the involvement of dopamine (DA) neurons in the substantia nigra pars compacta (SNc), traditionally associated with motor symptoms [10, 13–15]. Caudal raphe 5-HT neurons are essential for various autonomic processes, including the regulation of GI motility [16], which is often impaired in PD. Conversely, in the brain-first subtype, it is hypothesized that α-Syn pathology originates in the olfactory bulb and/or the cortico-limbic regions, with subsequent propagation to the brainstem and peripheral system [11]. It is also important to note that a dysfunctional 5-HT system is widely recognized as a significant risk factor for depressive disorder (DD) [17, 18]. Consistent with this, several reports indicate a positive correlation between reduced 5-HT neurotransmission and the severity of depressive and anxiety symptoms in PD [7, 19–21]. This is likely due to pathological alterations in the 5-HT system, as was recently reported in an α-synucleinopathy mouse model [22, 23], providing a biological substrate for the early manifestation of depression and GI dysmotility in PD.

Chronic inflammation is a key driver of the gut-brain pathology in PD. A compromised gut barrier can trigger systemic inflammation and immune responses that can exacerbate neuroinflammation and perpetuate the neurodegenerative process in PD. Indeed, recent studies have revealed a significant correlation between PD and inflammatory bowel diseases (IBD) [24–26], a group of chronic intestinal diseases that includes ulcerative colitis and Crohn’s disease. Similarly, a bidirectional association has been established between IBD and DD, whereby each condition can influence the development and course of the other. Patients with IBD are two to three times more likely to develop DD compared to the general population, and the prevalence of depressive symptoms in IBD patients is estimated to be between 21% and 27% [27, 28]. Furthermore, meta-analyses have demonstrated that individuals diagnosed with DD are at significantly higher risk of subsequently developing IBD [29, 30]. While the relationship between gut inflammation, leaky gut and PD/DD is becoming clearer [31, 32], the precise molecular regulators orchestrating these processes remain largely unknown.

An emerging yet under-explored shared pathophysiological mechanism involves abnormal microRNA (miRNA) expression. These small, non-coding RNAs regulate gene expression post-transcriptionally and are essential for preserving the integrity of both the intestinal epithelium and blood-brain barrier (BBB) [33–35]. For instance, while dysregulation of miRNAs such as miR-122a, miR-31, and miR-93 impairs gut barrier function, leading to gut inflammation and an increased susceptibility to gut-derived pathologies [36–38]; others, such as miR-126 and miR-132 are known to enhance BBB integrity [35, 39]. Accumulating evidence implicates dysregulated miRNAs as key regulators of immune-inflammatory pathways connecting gut inflammation with PD and DD [40, 41]. Based on this, we hypothesized that miRNAs reported to be differentially expressed in IBD would also be dysregulated in PD and DD. In this study, we analyzed postmortem human brain tissue and two established mouse models with PD and DD phenotypes. We identified three distinctive IBD-associated miRNAs, miR-199a-5p, miR-219a-5p, and miR-200a-3p, which were concordantly altered in the brains of patients with PD and DD, as well as in the mouse models. At the same time, we observed an increase in the expression of NFκB1 mRNA, a direct target of miR-199a-5p, and downstream inflammatory pathways. Our findings imply that a common, miRNA-mediated inflammatory mechanism may underlie the pathological triad of PD, DD and GI dysfunction.

## Materials and methods

### Postmortem human brain samples

All procedures were approved by the Ethics Committee of the Spanish National Research Council (Project Ref. PID2019-105136RB-I00) and were performed in compliance with the research and ethics guidelines for postmortem brain studies in Spain. Postmortem brain tissue from the dorsolateral prefrontal cortex (dlPFC) and caudate nucleus (CAU) was obtained during autopsy at the Basque Institute of Legal Medicine, Bilbao, Spain. The study included a total of 28 subjects with DD and 28 matched, non-psychiatric controls. The DD diagnoses were performed by a psychiatrist based on DSM-IV criteria and documented in the subjects’ medical records. Control subjects were confirmed to have no history of neurologic or psychiatric conditions, with their cause of death being mainly accidental. In contrast, 23 of the DD subjects died by suicide, while 5 died from natural causes. Each pair of subjects was matched for age, gender, postmortem interval (PMI), and storage time when possible. Following dissection, samples were stored at −80 °C until assays were performed. Full toxicological analyses were carried out on all subjects’ blood samples to screen for psychotropic drugs. Detailed cohort characteristics are presented in **Table 1**.

**Table 1.** Demographic and clinical characteristics of each group used in the present study.

| **Depressive Disorder (DD)** |  |  |  |  |  |  |  |  |  |
| --- | --- | --- | --- | --- | --- | --- | --- | --- | --- |
|  | No. of subjects | Gender |  | Age (years) |  | Postmortem interval (h) |  | AD drug | Cause of death |
|  |  | Male | Female | Male | Female | Male | Female |  |  |
| Control | 28 | 19 | 9 | 39.6 ± 13.9 | 42.1 ± 13.5 | 14.8 ± 1.9 | 18.9 ± 2.5 | 0 | Natural/accident |
| MDD | 28 | 19 | 9 | 39.8 ± 13.9 | 41.8 ± 13.6 | 20.6 ± 2.6 | 15.9 ± 2.7 | 7 | Natural/suicide |
| p-value | N/A | N/A | N/A | 0.954 | 0.959 | 0.084 | 0.433 | N/A | N/A |
| **Parkinson's disease (PD)** |  |  |  |  |  |  |  |  |  |
|  | No. of subjects | Gender |  | Age (years) |  | Postmortem interval (h) |  |  |  |
|  |  | Male | Female | Male | Female | Male | Female |  |  |
| Control | 10 | 3 | 7 | 74.7 ± 5.5 | 79.3 ± 3.7 | 11.1 ± 4.0 | 12.1 ± 1.9 |  |  |
| Braak 2-3 | 10 | 5 | 5 | 77.0 ± 3.9 | 88.0 ± 4.9 | 9.6 ± 1.7 | 8.8 ± 1.4 |  |  |
| Braak 5-6 | 10 | 5 | 5 | 78.4 ± 2.5 | 75.0 ± 3.7 | 10.1 ± 2.1 | 8.7 ± 2.8 |  |  |
| p-value | N/A | N/A | N/A | 0.736 | 0.939 | 0.833 | 0.460 |  |  |

Values denote mean ± SEM. N/A: not applicable. AD drug: antidepressant drug

Furthermore, in compliance with Spanish legislation and with the approval from local ethics committees, a cohort of 20 PD patients and 10 controls (male and female) was obtained from the Hospital Clinic-IDIBAPS biobank. Postmortem dlPFC and CAU samples were collected within a 3-to-20-hour interval after death and stored at −80 °C. The PD cases were pathologically classified as Braak stages 1–6 for Lewy body pathology, excluding those with tauopathies, vascular, or metabolic diseases. Controls were devoid of neurological, psychiatric, or metabolic disorders, showing no neuropathological issues besides sporadic Alzheimer’s. All PD patients, who had a disease duration of 8 to 18 years, had been treated for motor symptoms and typically died from infections, neoplasia, or acute heart disease. **Table 1** provides a comprehensive summary of subject details.

### Mice

Male and female C57BL/6J mice (10 weeks; n = 84 mice for the whole study, Charles River, Lyon, France) were housed under controlled conditions (22 ± 1°C; 12h light/dark cycle) with food and water available *ad-libitum*. Animal procedures were conducted in accordance with standard ethical guidelines (EU directive 2010/63 of 22 September 2010) and approved by the local ethical committee (University of Barcelona).

### Mouse model of PD with depressive-like phenotype (DD-PD)

In order to generate the DD-PD mouse model, a recombinant adeno-associated viral vector serotype-2/5 (AAV2/5) with chicken-β-actin (CBh) promoter and a shortened WPRE3 that encodes the human wild-type α-synuclein (AAV-h-α-Syn) was used to overexpress the transgene in raphe 5-HT neurons of male mice (dorsal raphe nucleus - DR), as previously reported [22, 23]. The AAV2/5 constructs were provided from The Michael J. Fox Foundation and produced and titered by the UNC Vector Core. Control group of animals received an empty vector (AAV-EV) with the same AAV2/5-CBh-WPRE3 backbone but containing non-coding stuffer DNA. For vector delivery, mice were anesthetized with isoflurane (4% induction, 2% maintenance) and placed in a stereotaxic frame. Mice were randomly infused with 1 μl of AAV-h-α-Syn or AAV-EV (concentration 0.3 x 10^13^ gc/mL) into raphe nuclei (anterior-posterior AP: -4.5; medial-lateral ML: -1.0; dorsal-ventral DV: -3.2 in mm, relative to bregma with an angle 20°), using a microinjector (KDS-310-PLUS, World Precision Instruments, Sarasota, FL, USA) at 0.4 µl/min rate. The needle was kept in place for an additional 5 min before slowly being withdrawn. All subsequent analyses were performed at 4 and 8 weeks post-infusion.

### Corticosterone-induced stress mouse model

We used the corticosterone (CORT) consumption-based stress model to generate a DD-like mouse model, as previously described [42, 43]. To do this, female mice were randomly distributed in control and CORT groups before the treatment, and animal weight balance was equally represented in both groups. CORT (Sigma-Aldrich) was dissolved in commercial mineral water and adjusted to a pH 7.0-7.4 with HCl. Decreasing CORT concentrations were presented to mice for 28 days: 30 μg/ml during 15 days (resulting in a dose of approximately 6.6 mg/kg/day), followed by 15 μg/ml (2.7 mg/kg/day) during 3 days, and 7.5 μg/ml (1.1 mg/kg/day) during 10 days for a gradual recovery of endogenous corticosterone plasma levels [42, 43]. CORT solution was available ad libitum in drinking water (opaque bottles) and was renewed every 72 hours. To verify CORT consumption, the weight of the bottles was checked each time the solution was renewed. Control mice followed the same experimental approach without CORT in their bottles. Mice from both groups were subjected to behavioral evaluation at day 36, and then were euthanized by cervical dislocation and brain samples were processed for qPCR analysis.

### Behavioral analysis

Different behavioral paradigms were used to evaluate motor, anxiety and depressive-like phenotype [22, 42]. All tests were performed between 10:00 and 16:00 h by an experimenter blind to mouse treatments. On the day of testing, mice were moved to a dimly lit behavioral room and left undisturbed for at least 1 h prior to testing. Mice were tested in random order and the equipment was thoroughly cleaned with ethanol 75% between each trial to minimize odors.

#### Open field test (OFT)

Motor activity was measured in a plexiglas open field box (35 × 35 × 40 cm) indirectly illuminated (25–40 lux) to avoid reflection and shadows. The box floor was covered with an interchangeable opaque plastic base that was replaced for each animal. Motor activity was recorded during 15 min by a camera connected to a computer (Video-track, Viewpoint, Lyon, France). The following variables were measured: horizontal locomotor and exploratory activity, defined as the total distance moved in cm including fast/large (speed > 10.5 cm/s) and slow/short movements (speed 3-10.5 cm/s), and the activation time including mean speed (cm/s) and resting time (s).

#### Tail suspension test (TST)

Mice were suspended 30 cm above the bench by adhesive tape placed approximately 1 cm from the tip of the tail. Sessions were videotaped for 6 min and the immobility time was measured (Smart, Panlab, Cornella, Spain).

#### Forced swim test (FST)

Mice were individually placed into a clear cylinder (15 cm diameter, 30 cm height) containing 20 cm of water maintained at 24–25°C and the time of immobility was measured for a total of 6 min test and represented in 2-min fractions. The test was recorded using a video camera system (Smart, Panlab, Cornella, Spain).

### Gut transit test

To measure total gut transit time, mice were administered 200 µL of a freshly prepared Carmine Red solution via a 21-gauge, round-tipped feeding needle. This oral gavage solution was prepared by dissolving Carmine Red (Sigma, C1022) to a final concentration of 6% in 0.5% methylcellulose (Sigma, M0512). Post-administration, mice were immediately housed in individual cages, and their stool was continuously monitored for the first appearance of the red-colored dye. The total gut transit time was recorded for each mouse as the interval between the oral administration of the dye and the excretion of the first red fecal pellet [44].

### RNA isolation

Total RNA was isolated from human postmortem dlPFC and CAU samples from patients with PD, DD, and control subjects, as well as from mouse medial prefrontal cortex (mPFC), caudate-putamen (CPu), and ileum tissues. The extraction was performed using the miRNeasy Mini Kit (Qiagen, Germany) following the manufacturer’s protocol. RNA concentration and purity were assessed by spectrophotometric analysis (NanoDrop™), with acceptable thresholds defined as 260/280 nm ≥ 1.8 and 260/230 nm ≥ 2.0. RNA integrity was assessed using an Agilent 2100 bioanalyzer, and only samples with an RNA Integrity Number (RIN) of 7 or higher were included in subsequent analyses.

### First-strand cDNA synthesis

For miRNA analysis, 2.5 µg of total RNA was first polyadenylated in a 50 µL reaction containing 1.5 U of E. coli poly(A) polymerase I (Thermo Fisher Scientific, AM2030) and 1 mM ATP (Thermo Fisher Scientific, R0441). The reaction was incubated at 37°C for 60 min. The polyadenylated RNA was then immediately reverse transcribed using the High-Capacity cDNA Reverse Transcription Kit (Thermo Fisher Scientific, 4368814) with a custom degenerate oligo-dT adapter primer (5′-GCACGCTACGACCTCAGAGTCACTATGGCACTTTTTTTTTTTTTTTTVN-3′, V = G/A/C and N = G/A/T/C). The reverse transcription was performed at 37°C for 120 min, followed by enzyme inactivation at 85°C for 5 min.

For mRNA analysis, 0.5 µg of total RNA was reverse transcribed using the High-Capacity cDNA Reverse Transcription Kit (Thermo-Fisher Scientific, 4368814). The reaction was performed according to the manufacturer’s protocol using the following thermal profile: 25°C for 10 min, 37°C for 120 min, and 85°C for 5 min.

### Analysis of miRNAs and mRNAs by qPCR

Expression levels of miRNAs (miR-199a-5p, miR-219a-5p, and miR-200a-3p) and mRNAs (tumor necrosis factor alpha - TNFα, interferon gamma - IFN-γ, interleukin (IL)-2, IL-6, nuclear factor kappa-light-chain-enhancer of activated B cells - NFκB1, ionized calcium binding adaptor molecule 1 - Iba1, glial fibrillary acidic protein – GFAP, and serotonin 2A receptor - 5-HT_2A_R) were quantified by SYBR Green-based quantitative PCR (qPCR) on an Applied Biosystems 7500 Real-Time PCR platform (Thermo Fisher). All reactions were performed in duplicate in a final volume of 10 µL, containing 1X PowerTrack™ SYBR™ Green Master Mix (Thermo Fisher), 300 nM of each primer, and 2.5 ng of cDNA template. The standard thermal protocol consisted of an initial enzyme activation at 95°C for 2 min, followed by 40 cycles of denaturation at 95°C for 15 s and an annealing/extension step at 60°C for 60 s. A melting curve analysis was performed after amplification to verify product specificity. Relative expression for all targets was calculated using the 2-ΔΔCT method. Quantification of each target miRNA was performed using a unique primer pair. Each pair consisted of a miRNA-specific forward primer and a specific reverse primer designed to be complementary to the oligo-dT adapter sequence used during cDNA synthesis. Expression levels were normalized to the endogenous control RNU6. For the quantification of target mRNAs, gene-specific forward and reverse primer pairs were used for each transcript. For data normalization, the endogenous housekeeping gene beta-actin (*ACTB*) was used as a reference. All primer sequences are listed in **Supplementary Table 1**.

### In silico miRNA target gene prediction

To identify biologically relevant pathways associated with the three IBD-associated miRNAs, a multi-algorithmic approach was implemented by integrating results from seven bioinformatic tools: miRWalk, MicroT, mirabel, miRDB, miRTargetLink, TargetScan, and MiRanda. We retained targets that were either experimentally validated or had a binding probability > 0.90 in the 3’ untranslated region (UTR), 5’ UTR, or coding DNA sequence (CDS) regions. Strict reproducibility thresholds were applied and only mRNA targets predicted by ≥5 or more independent algorithms were retained for downstream ontological profiling, thereby minimizing false-positive associations.

### Prediction-based functional pathways and gene ontology analysis

To functionally annotate the consensus list of miRNA target genes, an Over-Representation Analysis (ORA) was performed using the web-based GEne SeT AnaLysis Toolkit (WebGestalt; http://www.webgestalt.org). This analysis was conducted to identify enriched KEGG pathways and Gene Ontology (GO) terms, focusing specifically on the Biological Process domain. The human genome (GRCh38) was used as the reference set, and statistical significance was determined using a hypergeometric test with a Benjamini-Hochberg false discovery rate correction (FDR < 0.05). KEGG pathways and GO terms, were ranked according to their enrichment scores. Only pathways containing at least five of our target genes were retained.

To prioritize neurobiological relevant pathways, enriched terms were manually curated to retain those associated with inflammation, synaptic transmission, and nervous system development and function. Redundant or hierarchical parent/child GO terms were consolidated to avoid over-representation. To represent the relationships between the most significant KEGG pathways, interaction networks were generated using Cytoscape, where nodes represented functional terms scaled by gene set size, and edge thickness are proportional to the degree of gene overlap. Enriched terms were cross-referenced with published literature on MDD, PD and IBD pathology to ensure biological relevance.

### Immunofluorescence

Mice were anaesthetized with pentobarbital and transcardially perfused with 4% PFA in sodium-phosphate buffer (pH 7.4). Brains were extracted, post-fixed 24 h at 4°C in the same solution, and placed in gradient sucrose solution 10–30% for 3 days at 4°C. After cryopreservation, serial 30 μm-thick sections were cut to obtain raphe nuclei, mPFC, and CPu. Immunofluorescence procedure was performed for h-α-Syn (anti-h-α-Synuclein clone Syn 211, 1:2000; ref.: AHB0261, Thermo Fisher Scientific, Waltham, MA, USA), mouse/human-α-Syn (anti-m/h-α-Syn, 1:500; ref.: ab212184, Abcam, Cambridge, UK), tryptophan hydroxylase TPH (anti-TPH, 1:2500; ref.: AB1541, Sigma-Aldrich, Madrid, Spain), serotonin transporter SERT (anti-SERT, 1:2500; ref.: 24330, Immunostar, Hudson, WI, USA), glial fibrillary acidic protein GFAP (anti-GFAP, 1:1000; ref.: G3893, Sigma-Aldrich, Madrid, Spain) and ionized calcium-binder adapter molecule 1 Iba1 (anti-Iba1, 1:1000; ref.: 019-197741, Fujifilm Wako Chemicals Europe, Fuggerstraße, Neuss, Germany). Sections were washed in 1x PBS (pH 7.4) and 1x PBS/Triton 0.2%, and incubated in blocking solution (1x PBS/Triton containing 0.02% gelatin and the corresponding normal serum for the secondary antibody host) for 2 hours at room temperature. Primary antibody was incubated overnight at 4°C, followed by incubation with the respective secondary antibodies for 2 hours at room temperature (**Supplementary Table 2**). Finally, nuclei were stained with Hoestch dye (1:10.000; ref.: H3570, Life Technologies, Carlsbad, CA, USA) for 10 min and the sections were mounted in Entellan (Electron Microscopy Sciences, Hatfield, PA, USA).

### Confocal fluorescence microscopy

The intracellular localization of h-α-Syn in TPH-positive cells in the DR, and the intracellular h-α-Syn distribution in 5-HT axons were examined by confocal microscopy using an inverted Nikon Eclipse Ti2-E microscope (Nikon Instruments, Tokyo, Japan) attached to the spinning disk unit Andor Dragonfly 200 (Oxford Instruments Company, Abindong, UK). For all experiments, a Plan Apochromatic 10-20x, numerical aperture (NA) 0.45 objective was used. A high-precision motorized stage was used to collect the large-scale 3D mosaics of each tissue section. Individual image tiles were 2048 × 2048 pixels with a z-section of 35 µm. For specific regions, we used an oil-immersion objective (Plan Apochromatic Lambda blue 40x, NA 1.4). Samples were excited with 405, 488, and 561 nm laser diodes, respectively. The beam was coupled into a multimode fiber going through the Andor Borealis unit reshaping the beam from a Gaussian profile to a homogenous flat top, and from there it was passed through the 40 µm pinhole disk. Tissue sections were imaged on a high resolution scientific complementary metal oxide semiconductor (sCMOS) camera (Zyla 4.2, 2.0 Andor, Oxford Instruments Company).

Fusion software (Andor, Oxford Instruments Company) was used for acquisition and for image processing before analysis. Image stitching and deconvolution were performed using Fusion software (Andor, Oxford Instruments Company). Image analysis was performed with Image J/Fiji open source software (Wayne Rasband, NIH, Bethesda, MD, USA) using ImageJ Macro Language to develop custom macros. Briefly, maximum projections of the channels of interest (red and green) were obtained for each section, and regions of interest (ROI) were delimited for each brain section to cover the maximum analyzable area. Quantification was performed using a group size of six animals and three coronal sections per animal. ROIs were systematically defined in all animals and brain regions. One ROI was analyzed per section in the DR, while two ROIs were analyzed per section (bilaterally) in the brain projection areas. These sampling variables were kept constant for all animals in each experimental group.

Since the number of GFAP/Iba1-labelled cells could not be directly counted due to their irregular morphology and heterogeneous distribution, automated macro scripts were developed for quantification. Before running these automated macro scripts, images were segmented using the MaxEntropy automatic threshold and processed with various filters to reduce background noise and remove outliers. For the analysis of GFAP-labelled cells, the Skeletonize plugin in ImageJ was applied following the protocol established by Marques et al [45]. This process transformed astrocytes into branching structures, allowing for their independent quantification. The results obtained with Analyze Skeleton were filtered to exclude structures with fewer than two branches, after which the number of branched structures was counted. Regarding Iba1-labelled cells, we used the Microglia Morphometry plugin in ImageJ, developed by Martínez et al [46]. Microglial cells were identified and labeled as independent particles. Subsequently, the Analyze Particles function generated a results table indicating the number of cells and the area of each particle, allowing for the determination of the mean cell area.

### Imaris 3D surface rendering

In order to generate 3D surface representations of the structures of interest, confocal images were acquired for the channels of interest (red and green) using a 60x objective. The z-step size was optimized to maximize the number of planes and enhance image quality. The 3D reconstructions were performed using Imaris Image Analysis Software Version 10.2.0 (Bitplane, Zurich, Switzerland). Briefly, surfaces were generated for each channel separately using different settings within the Surface tool, adjusting the parameters to match the specific labelling characteristics of each protein. For cellular markers, including TPH, Iba1 and m/h-α-Syn, we used the Machine Learning-based method, in which the program was trained iteratively by manually classifying regions of the image as background or foreground, thereby improving segmentation accuracy. Afterwards, the software segmented the surfaces into individual cells using a defined minimum particle diameter (7 μm in our case). Finally, a size filter was applied to retain only the particles corresponding to the expected cell size and morphology.

For h-α-Syn staining, which shows a more diffuse protein distribution, as well as for GFAP and SERT-labelled images, which exhibit an irregular fibrillar morphology, surfaces were generated manually using the Absolute Intensity Threshold option. After adjusting the threshold to optimize rendering and achieve clear separation from the background, particles were segmented by diameter (1 μm for h-α-Syn and 7 μm for GFAP, no segmentation was required for SERT staining) and the size filter was subsequently applied to retain only the objects matching the expected size and morphology.

### Statistical analyses

All values are the mean ± standard error of the mean (SEM). Data from individual human and mouse samples were represented by individual points. Statistical comparisons are performed using GraphPad Prism 10.4.1 software. Outlier values were identified and excluded using the ROUT test (i.e., Extreme Student Zed Deviate – ESD method). When appropriate, Student’s *t*-test, one-way and two-way ANOVA analysis followed by a Bonferroni *post hoc* test were performed, as indicated in the results and figure legends. The non-parametric Kruskal-Wallis test followed by the Dunnett’s multiple comparison test was used to analyze differences in means between the control and the different stages of PD and in DD. Differences were considered significant if p < 0.05. A summary of the statistical analysis is shown in the **Supplementary Excel File**.

## Results

### Identification of common dysregulated miRNAs linked to IBD in postmortem brains of PD and DD

Previous sequencing data revealed changes in miRNA expression in patients with IBD, such as the downregulation of miR-199a-5p, miR-219a-5p, and the upregulation of miR-200a-3p playing a significant role in affecting intestinal permeability, inducing visceral hypersensitivity, and aggravating inflammation [40, 47]. Particularly, miR-200a-3p inhibits SERT mRNA expression [48], leading to increased gut 5-HT levels, which are positively correlated with IBD susceptibility [49]. Given the growing evidence linking gut-brain axis disruption to systemic and central inflammation in neurodegenerative and neuropsychiatric disorders, we examined whether these IBD-associated miRNAs also exhibit changes in postmortem brain tissue samples from patients with PD and DD.

We first quantified the levels of miR-199a-5p, miR-219a-5p, and miR-200a-3p in the dlPFC and CAU of subjects with PD at different stages (**Fig. 1A-F**). In the dlPFC, we detected a significant downregulation of miR-199a-5p at both early (Braak 2-3, B2-3) and late (Braak 5-6, B5-6) stages of PD (p < 0.01) compared to the control group (**Fig. 1A**). Similarly, miR-219a-5p levels were reduced, reaching statistical significance at the late B5-6 stage (p < 0.01) (**Fig. 1B**). In contrast, miR-200a-3p was significantly upregulated in the dlPFC of PD subjects at both B2-3 (p < 0.01) and B5-6 (p < 0.05) stages compared to the controls (**Fig. 1C**). A similar pattern of dysregulation was found in the CAU of patients with PD. Levels of miR-199a-5p and miR-219a-5p were significantly reduced at both early and late PD stages (p < 0.01) compared to the control subjects (**Fig. 1D, E**). On the contrary, miR-200a-3p expression was increased in CAU of PD at B2-3 (p < 0.01) and B5-6 (marginal effect p = 0.0560) compared to the control subjects (**Fig. 1F**).

**Figure 1.**
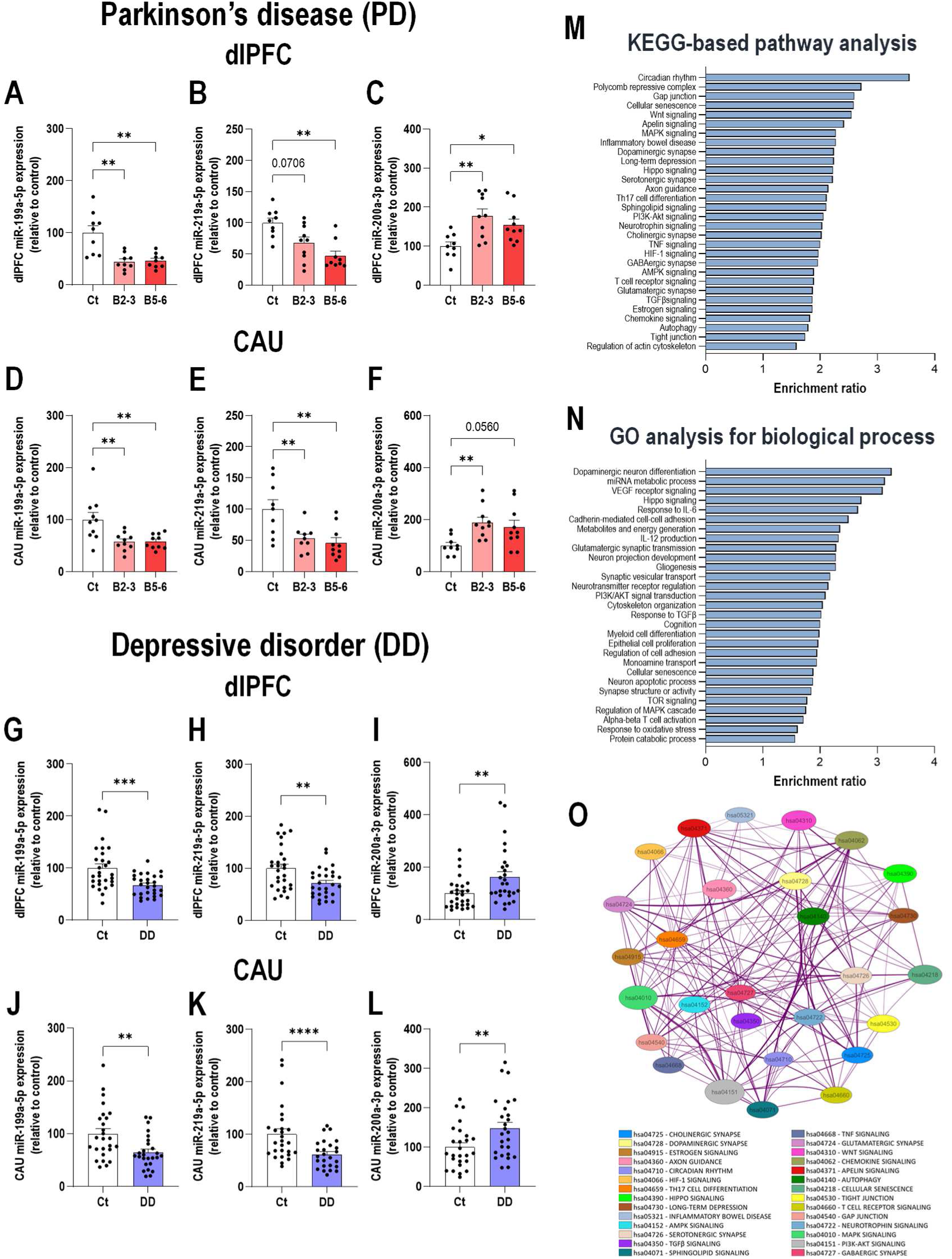
Identification of dysregulated miRNAs in postmortem brains samples of patients with Parkinson’s disease (PD) and depressive disorder (DD). **A-C)** Expression levels of miR-199a-5p, miR-219a-5p, and miR-200a-3p in the dorsolateral prefrontal cortex (dlPFC) of patients with PD at the early Braak 2-3 (B2-3) and late Braak 5-6 (B5-6) stages compared to the control group (Ct). **D-F)** Expression levels of the three miRNAs in the caudate nucleus (CAU) of the same patients with PD compared to the Ct group. **G-I)** Expression levels of miR-199a-5p, miR-219a-5p, and miR-200a-3p in the dlPFC of patients with DD compared to the respective Ct group. **J-L)** Expression levels of the three miRNAs in the CAU of the same patients with DD compared to the Ct group. **M)** KEGG-based pathway analysis of the predicted target genes of the dysregulated miRNAs. The bar chart shows the enrichment ratio of the most significantly enriched pathways. **N)** Gene Ontology (GO) analysis for the biological process category. The bar chart displays the enrichment ratio of the most significant processes associated with the predicted miRNA target genes. **O)** Enrichment network visualizing the relationships among the identified KEGG functional pathways. Each node represents an enriched term, color-coded by functional cluster. The connecting edges indicate the degree of overlap based on shared genes between the terms. Data is plotted as mean ± SEM. Data points represent individual human samples. * p < 0.05, ** p < 0.01, *** p < 0.001, **** p < 0.0001 versus controls (detailed statistical analysis in supplementary excel file).

Next, we analyzed the levels of these miRNAs in postmortem dlPFC and CAU samples from patients with DD and healthy subjects (**Fig. 1G-L)**. We found similar expression patterns, with reductions in miR-199a-5p (dlPFC, p < 0.001; CAU, p < 0.01; **Fig. 1G, 1J**) and miR-219a-5p (dlPFC, p < 0.01; CAU, p < 0.0001; **Fig. 1H, 1K**), and increased expression of miR-200a-3p (dlPFC, p < 0.01; CAU, p < 0.01; **Fig. 1I, 1L**) in the postmortem samples from patients with DD compared to the control group. Additional analyses considering the sex variable showed a similar profile of negative regulation of miR-199a-5p and miR-219a-5p, and positive regulation of miR-200a-3p in the dlPFC and CAU of male and female patients with DD compared with the respective control group (**Suppl Fig. 1)**.

Following the significant dysregulation of miR-199a-5p and miR-219a-5p (downregulated) and miR-200a-3p (upregulated) in the dlPFC and CAU of PD and DD patients, we performed a functional enrichment analysis on their predicted target genes (**Fig. 1M, 1O**). The results reveal that these miRNAs converge on a core set of interconnected pathways implicated in both PD and DD pathology, particularly those governing neurotransmitter systems, neuroinflammation, and synaptic plasticity. The analysis of KEGG pathways highlighted a profound disruption of monoamine signaling. The dopaminergic synapse pathway (KEGG: hsa04728, FDR < 0.01) was significantly enriched, reflecting a direct impact on the system primarily affected in PD. Furthermore, while a specific serotonergic pathway was not a top hit, key genes involved in 5-HT signaling, including the *HTR2A* and *HTR2C*, were integral components of other highly significant pathways, including PI3K-Akt signaling (KEGG: hsa04151, FDR < 10^-5^), indicating a broader dysregulation of monoaminergic neurotransmission (**Fig. 1M, 1O**).

Crucially, this disruption is linked to a strong neuroinflammatory and metabolic stress signature. Core inflammatory cascades, including MAPK signaling (KEGG: hsa04010, FDR < 10^-5^) and the aforementioned PI3K-Akt signaling (KEEG: hsa04151, FDR < 10⁻^5^), were among the most enriched pathways. These pathways are not only central to microglial activation and cytokine response in the brain but are also well-established drivers of systemic inflammation, such as in IBD (KEEG: hsa05321, FDR < 0.05), providing a molecular link for the gut-brain axis (**Fig. 1M, 1O**).

This convergence of altered neurotransmission and inflammation appears to precipitate a widespread disruption of synaptic architecture and function (**Fig. 1N, 1O**). The Gene Ontology (GO) analysis was dominated by terms related to synaptic biology, including neuron projection development (GO: 0010975, FDR < 1×10⁻^9^), regulation of synapse structure or activity (GO: 0050803, FDR < 0.01), and synaptic vesicle transport (GO: 0099003, FDR < 10^-4^). Highly significant enrichments were found in processes such as branching structure morphogenesis (GO: 0001763, FDR < 10⁻^8^), dopaminergic neuron differentiation (GO: 0071542, FDR < 0.01), and axonogenesis (GO: 0007409, FDR < 10⁻^8^), as well as in cognition (GO: 0050890, FDR < 10^-4^). This suggests that the primary pathological consequence of this altered miRNA profile is a failure to maintain synaptic integrity, likely driven by the combined deficits in neurotransmitter signaling and the chronic inflammatory environment. Intriguingly, miRNA metabolic process (GO: 0010586, FDR < 10^-5^) was also highly enriched, suggesting a potential auto-regulatory loop where these miRNAs may influence their own biogenesis, thus perpetuating the pathological state (**Fig. 1N, 1O**).

In parallel, a strong association with cellular stress responses was observed. Pathways like response to oxidative stress (GO: 0006979, FDR < 0.01), cellular senescence (KEGG: hsa04218, FDR < 10⁻^4^), and HIF-1 signaling (KEGG: hsa04066, FDR < 0.05) were significantly enriched (**Fig. 1M-O**). The involvement of key genes like *SIRT1*, *HIF1A*, and *TP53* suggests that this miRNA profile compromises the ability of neurons to manage metabolic and oxidative stress, thereby accelerating neurodegenerative processes. Finally, the results directly link these miRNA dysregulations to neuroinflammation. Pathways such as response to IL-6 (GO: 0070741, FDR < 0.05) and canonical NFκB1 signaling (GO: 0007249, FDR < 0.05), along with TNF signaling (KEGG: hsa04668, FDR < 0.05) were affected. This suggests that the altered miRNAs modulate the immune response in the brain. Concurrently, an impact on structural integrity was identified through pathways like adherent junction (KEGG: hsa04520, FDR < 0.001) and focal adhesion (KEGG: hsa04510, FDR < 0.05), which could reflect a dysfunction in cell-cell and cell-matrix interactions in the affected brain tissue (**Fig. 1M-O**). In summary, the dysregulation of miR-199-a5p, miR-219a-5p, and miR-200a-3p in the PD and DD brain appears to orchestrate a coordinated breakdown of multiple cellular defense and maintenance systems. Their convergent impact on neurotrophic and developmental signaling cascades (Wnt, PI3K-Akt), synaptic and axonal function, cellular stress management, and the neuroinflammatory response establishes these miRNAs as key contributors to the pathogenesis of PD and DD. Their simultaneous action on monoamine neurotransmission, neuroinflammatory signaling, and synaptic plasticity provides a solid molecular basis that could partly explain the clinical comorbidity between PD and DD.

### Altered inflammatory response markers in postmortem brain samples from patients with PD and DD

In order to establish whether the dysregulation of miR-199a-5p and miR-219a-5p (downregulated) and miR-200a-3p (upregulated) is associated with central neuroinflammatory processes, we systematically analyzed the mRNA levels of key cytokines such as TNFα, IFN-γ, IL-2, IL-6, as well as the expression of the transcription factor NFκB1 in postmortem brain samples from patients with PD and DD.

In the dlPFC of PD patients, we detected significantly increased mRNA levels of TNFα and IFN-γ at the early (B2-3, p < 0.0001 and p < 0.05, respectively) and late (B5-6, p < 0.05) stages, respectively (**Fig. 2A, B**). Additionally, IL-2 (p < 0.01) and IL-6 (p < 0.001) expression was enhanced in the dlPFC of PD patients at the B5-6 stage (**Fig. 2C, 2D**). Furthermore, a significant increase (p < 0.05) in NFκB1 expression was detected in the dlPFC of PD patients at B2-3 and B5-6 compared to the control group (**Fig. 2E**). A similar neuroinflammatory profile was found when analyzing these same markers in the CAU of PD patients. Specifically, we detected elevated levels of TNFα in the early stage (p < 0.01) and IFN-γ in the late stage (p < 0.05) (**Fig. 2F, 2G**), with no changes observed in IL-2 and IL-6 expression (**Fig. 2H, 2I)**. As in the dlPFC, NFκB1 expression was marginally or significantly increased in the CAU of PD patients at B2-3 (p = 0.0695) and B5-6 (p < 0.05) compared to the control group (**Fig. 2J**).

**Figure 2.**
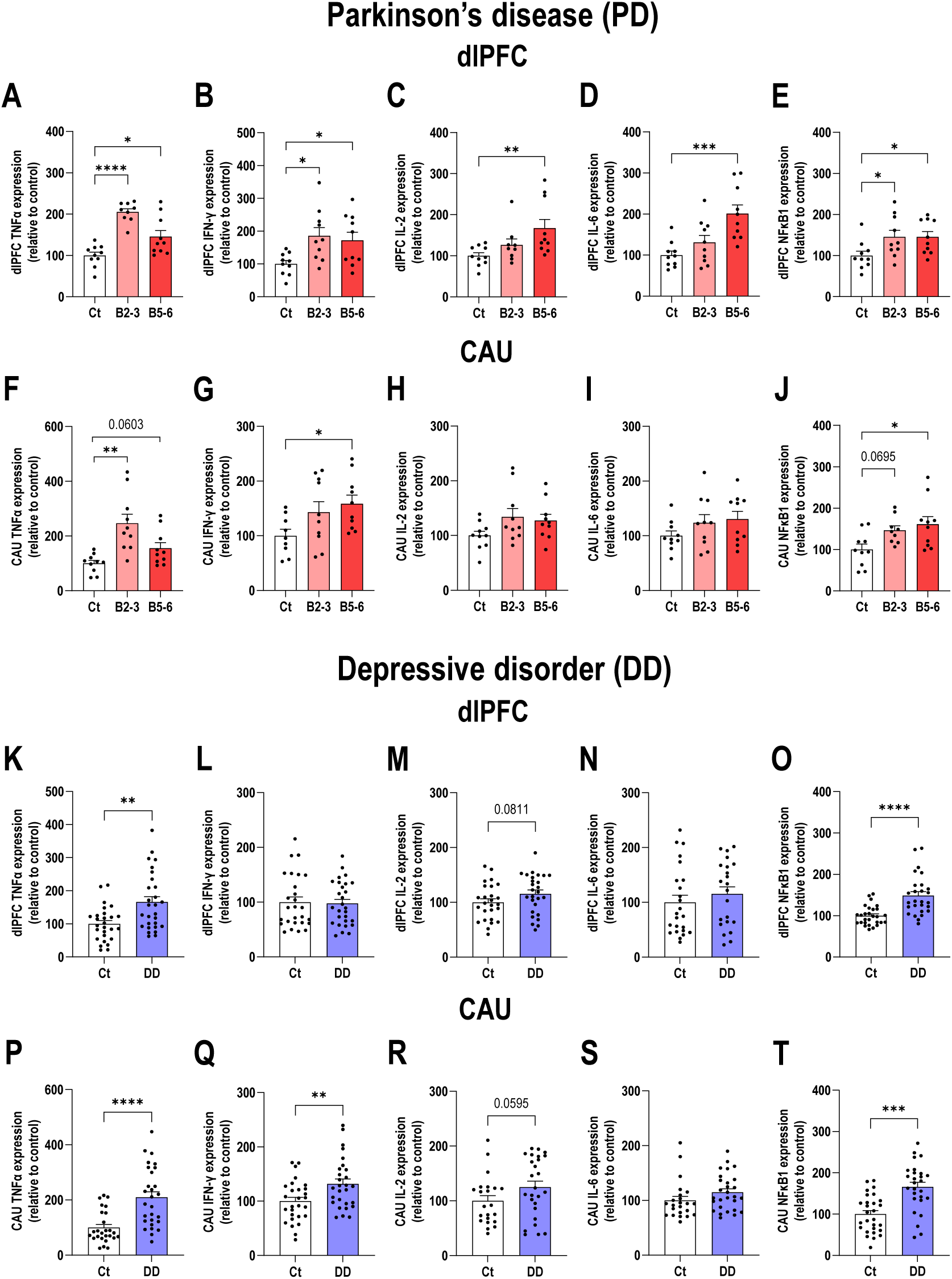
Changes in inflammatory response markers in postmortem brains samples of patients with Parkinson’s disease (PD) and depressive disorder (DD). **A-E)** Expression levels of TNFα, IFN-γ, IL-2, IL-6, and NFκB1 mRNAs in the dorsolateral prefrontal cortex (dlPFC) of patients with PD at the early Braak 2-3 (B2-3) and late Braak 5-6 (B5-6) stages compared to the control group (Ct). **F-J)** Expression levels of these inflammatory markers in the caudate nucleus (CAU) of the same patients with PD compared to the Ct group. **K-O)** Expression levels of TNFα, IFN-γ, IL-2, IL-6, and NFκB1 mRNAs in the dlPFC of patients with DD compared to the respective Ct group. **P-T)** Expression levels of these inflammatory markers in the CAU of the same patients with DD compared to the Ct group. Data is plotted as mean ± SEM. Data points represent individual human samples. * p < 0.05, ** p < 0.01, *** p < 0.001, **** p < 0.0001 versus controls (detailed statistical analysis in supplementary excel file).

Next, we analyzed the cytokine profile in the dlPFC and CAU of patients with DD. We observed increased expression of TNFα in the dlPFC (p < 0.01, **Fig. 2K**) and CAU, p < 0.0001; **Fig. 2P**), as well as increased expression of IFN-γ in the CAU only (p < 0.01; **Fig. 2L, 2Q**). There were also marginal effects in IL-2 expression in both the dlPFC (p = 0.0811; **Fig. 2M**) and the CAU (p = 0.0595; **Fig. 2R**), but no changes were observed in IL-6 expression (**Fig. 2N, 2S**). Furthermore, elevated levels in NFκB1 expression were detected in the dlPFC (p < 0.0001, **Fig. 2O**) and CAU (p < 0.001, **Fig. 2T**) compared to the control group. We performed an additional analysis by sex in the DD and control cohorts. The increase in TNFα and NFκB1 levels was significant in both the dlPFC and CAU in male and female patients with DD compared to the respective control group. Similarly, the increase in IFN-γ levels remained in the CAU when the analysis was based on sex. The sex-based analysis also revealed a significant increase in IL-2 expression in the dlPFC and CAU in DD males and only in the CAU in DD females compared to the respective controls. In particular, we detected a significant increase in IL-6 levels in the CAU of females with DD compared to respective control group (**Suppl Fig. 2A-T**).

To further investigate the cellular nature of the inflammatory response observed, we analyzed the expression of Iba1 and GFAP as indicators of microglial and astroglial activation. We found increased Iba1 and GFAP levels in the dlPFC (p < 0.05) and CAU (p < 0.05) in PD patients at late B5-6 stage (**Fig. 3A, 3B, 3D, 3E**). Marginal effects were observed in GFAP levels in the dlPFC (p = 0.0596) and Iba1 expression in the CAU (p = 0.0677) in the early B2-3 stage compared to the control group (**Fig. 3B, 3D**). Similarly, Iba1 levels were increased in both brain areas (dlPFC, p < 0.001; CAU, p < 0.01) in DD patients compared to controls (**Fig. 3G, 3J, Suppl Fig. 3**). However, GFAP expression was reduced in the dlPFC in DD patients (p < 0.05; **Fig. 3H**), but was not altered in CAU (**Fig. 3K**) compared to controls. Marginal effects on GFAP expression were detected after sex-based analyses (**Suppl Fig. 3**).

**Figure 3.**
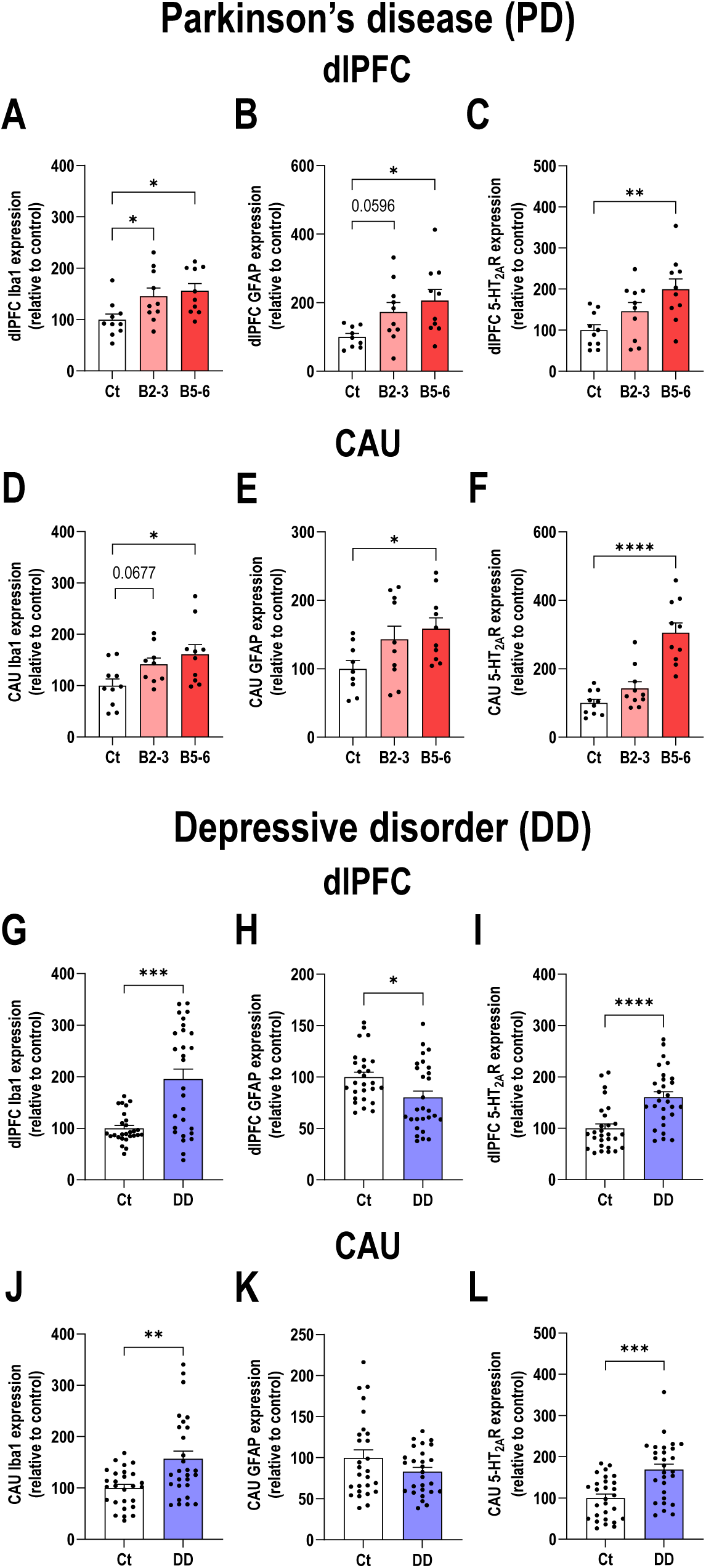
Changes in glial and serotonin (5-HT) signaling markers in postmortem brains samples of patients with Parkinson’s disease (PD) and depressive disorder (DD). **A-C)** Expression levels of Iba1, GFAP, and 5-HT_2A_R mRNAs in the dorsolateral prefrontal cortex (dlPFC) of patients with PD at the early Braak 2-3 (B2-3) and late Braak 5-6 (B5-6) stages compared to the control group (Ct). **D-F)** Levels of expression of these cellular markers in the caudate nucleus (CAU) of the same patients with PD compared to the Ct group. **G-I)** Expression levels of Iba1, GFAP, and 5-HT_2A_R mRNAs in the dlPFC of patients with DD compared to the respective Ct group. **J-L)** Levels of expression of these cellular markers in the CAU of the same patients with DD compared to the Ct group. Data is plotted as mean ± SEM. Data points represent individual human samples. * p < 0.05, ** p < 0.01, *** p < 0.001, **** p < 0.0001 versus controls (detailed statistical analysis in supplementary excel file).

Given that (i) analysis of the predicted target genes associated with the evaluated miRNAs impliates the *HTR2A*/*HTR2C* genes in 5-HT signaling and (ii) the relationship between the 5-HT system and the comorbidity of PD and DD [8, 15], we also evaluated the 5-HT_2A_R mRNA expression in brain samples from patients with PD and DD. The analysis showed increased 5-HT_2A_R expression in the dlPFC (p < 0.01; **Fig 3C**) and CAU (p < 0.0001, **Fig 3F**) in late-stage PD compared to controls. Similarly, we observed sex-dependent increases in 5-HT_2A_R expression in the dlPFC (p < 0.0001; **Fig 3I, Suppl Fig. 3)** and CAU (p < 0.001; **Fig 3L, Suppl Fig. 3**) in DD patients compared to controls.

### Pathological overexpression of α-Syn in raphe 5-HT neurons in mice triggers a global glial response, a depressive phenotype and GI dysfunction

We have recently developed a mouse model of PD displaying depressive phenotype (DD-PD), which is associated with oligomeric α-Syn and early dysfunction at the synaptic and brain network levels. The pathological accumulation and aggregation of α-Syn in the 5-HT system, induced by the local infusion of AA2/5-CBh-WPRE3A vector (denoted as AAV-α-Syn) in the DR, results in progressive α-synucleinopathy and the functional disconnection of brain circuits that are crucial for emotional regulation, as analyzed by fMRI [23]. Here, we used the same DD-PD-like mouse model to extend these observations, confirming the accumulation of human α-Syn protein in 5-HT TPH-positive cells 4 and 8 weeks later using in-depth analysis and visualization of 3D Imaris confocal images (**Fig. 4A, 4B**). Next, we confirmed the abundant and progressive presence of human α-Syn-positive fibers in efferent 5-HT brain regions, such as the mPFC and CPu (**Fig. 4C, 4D**), as previously reported [22, 23].

**Figure 4.**
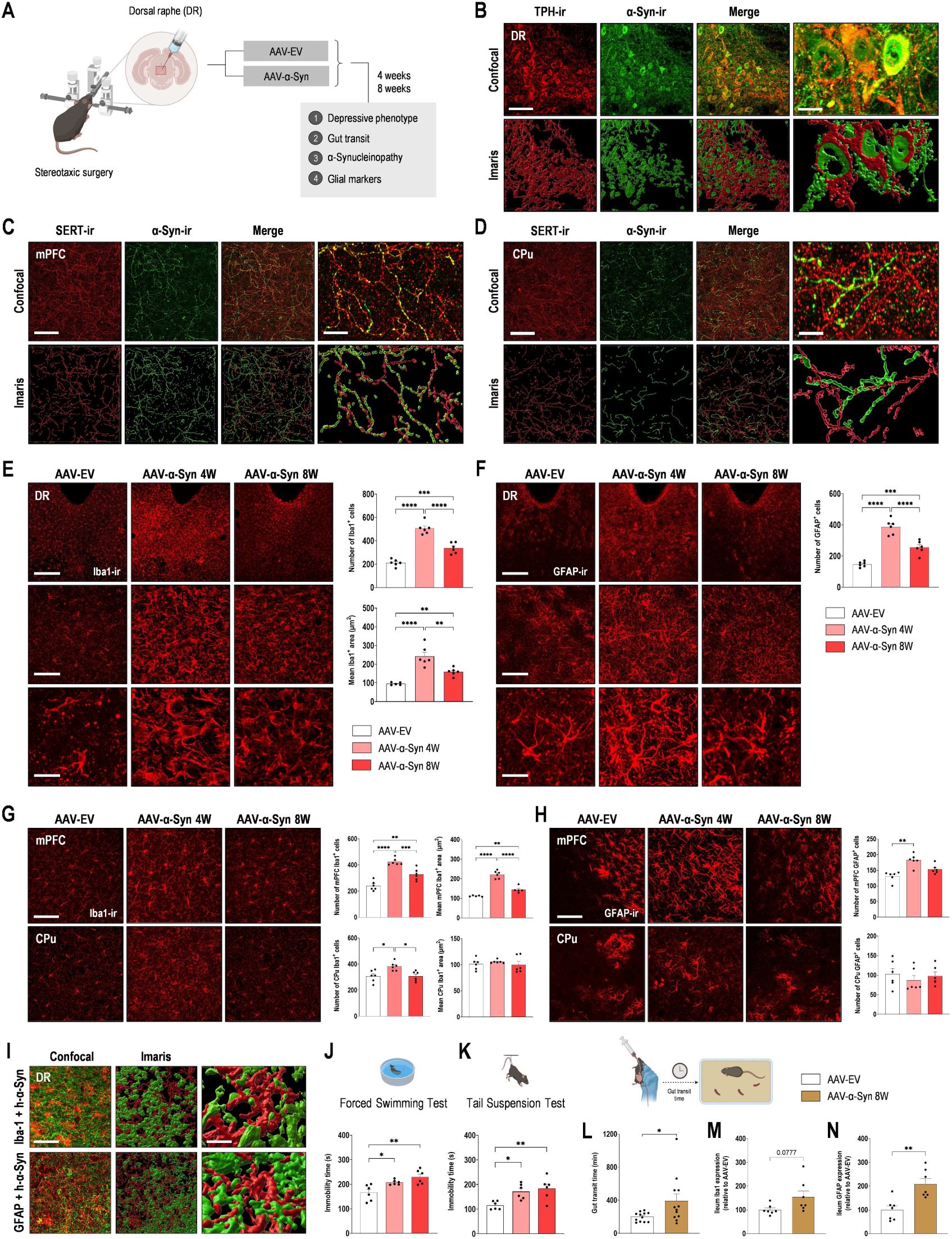
Overexpression of human α-Syn in raphe serotonin (5-HT) neurons in mice. **A)** Experimental design. Male mice received 1 μl of AAV2/5-CBh-WPRE3 construct to drive expression of human α-Syn (AAV-α-Syn) or empty AAV2/5-CBh-WPRE3 vector containing non-coding (null) stuffer DNA (AAV-EV) into dorsal raphe nucleus (DR) and were euthanized at 4 and 8 weeks (W) post-infusion [23]. **B)** Representative confocal images and Imaris 3D reconstructions showing the co-localization of human α-Syn protein and tryptophan hydroxylase-TPH^+^ cells in the DR of mice after AAV2/5 injection. Scale bar: 50 μm (low) and 10 μm (high). **C-D)** Representative confocal images and Imaris 3D reconstructions showing co-localization of human α-Syn^+^/serotonin transporter-SERT^+^ fibers in 5-HT projection brain regions, such as medial prefrontal cortex (mPFC), and caudate putamen (CPu) of mice. Scale bar: 25 μm (low) and 10 μm (high). **E)** Representative confocal images of Iba1^+^ microglia. Scale bar: 200, 50, and 10 μm, respectively. Bar chart showing a significant increase in the number and size of Iba1^+^ microglia in the DR at 4 and 8 weeks post-infusion. **F)** Representative confocal images of GFAP^+^ astrocytes. Scale bar: 200, 50, and 10 μm, respectively. Bar chart showing a significant increase in the number GFAP^+^ astrocytes in the DR at 4 and 8 weeks post-infusion. **G)** Representative confocal images and quantification of Iba1^+^ microglia in the mPFC and CPu. Scale bar: 50 μm. **H)** Representative images and quantification of GFAP^+^ astrocytes in the mPFC and CPu. Scale bar: 50 μm. **I)** High-magnification confocal images and Imaris 3D reconstructions showing the co-localization of human α-Syn^+^ (red) within Iba1^+^ microglia (green) or GFAP^+^ astrocytes (green) in the DR at 8 weeks post-infusion. Scale bar: 50 μm (low) and 10 μm (high). **J-K)** Mice with human α-Syn accumulation displayed depressive-like behaviors, including increased immobility time in the forced swimming and tail suspension tests. **L)** Whole gut transit quantification at 8 weeks post-infusion. **M)** Expression level of Iba1 mRNA in the ileum. **N)** Expression level of GFAP mRNA in the ileum. Data points represent individual mice. * p < 0.05, ** p < 0.01, *** p < 0.001, **** p < 0.0001 versus controls (detailed statistical analysis in supplementary excel file).

Since aggregated α-Syn activates microglia, and in turn, activated microglia are involved in both the clearance of α-Syn aggregates and the generation of pro-inflammatory responses that may be detrimental to neuronal health [50, 51], we then determined the density of the microglial marker Iba1 in the DR, mPFC and CPu in the DD-PD mouse model (**Fig. 4E, 4G**). The evaluation of Iba1-ir in the DR revealed a significant increase in reactive microglia in the DD-PD mouse model, which was greater at 4 weeks than at 8 weeks after AAV infusion (**Fig. 4E**). Analysis of confocal microscopy images showed an increase in both the number of Iba1-positive cells (4 weeks, p < 0.0001; 8 weeks, p < 0.001) and the area occupied by Iba1-positive microglia (4 weeks, p < 0.0001; 8 weeks, p < 0.01) in the DD-PD mice compared to the control mice. In addition, we also found significant increases in the number of Iba1-positive cells in the mPFC and CPu of DD-PD mice compared to the control group (**Fig. 4G**). Similar to DR, the effects on the Iba1-ir signal were greater in DD-PD mice 4 weeks later compared to 8 weeks (4 weeks: mPFC, p < 0.0001 and CPu, p < 0.05; 8 weeks: mPFC, p < 0.01). A significant increase in the area occupied by Iba1-positive microglia was only detected in the mPFC of the DD-PD group compared to the control group at both 4 and 8 weeks (p < 0.0001 and p < 0.01, respectively; **Fig. 4G**).

Reactive astrogliosis were also analyzed in the DR, mPFC, and CPu from all groups (**Fig. 4F, 4H**). Although no statistical differences were observed in the number of GFAP-positive astrocytes in the CPu in control and DD-PD mice, GFAP-ir reactive astroglia levels were statistically higher in the DR in DD-PD mice compared to control group at 4 weeks (p < 0.0001) and 8 weeks (p < 0.001) post-AAV infusion (**Fig. 4F, 4H**). In mPFC, we found increased number of GFAP-positive astrocytes only at 4 weeks post-AAV infusion (p < 0.01; **Fig. 4H**). Furthermore, we also found Iba1-positive and GFAP-positive cells contacting with human α-Syn, mainly in the DR of DD-PD mice, as seen in confocal 3D images (**Fig. 4I**).

We then confirmed that DD-PD mice exhibited a depressive-like behavior, spending more time immobile in the FST and TST at 4 (p < 0.05) and 8 weeks (p < 0.01) than the control group (**Fig. 4J, 4K**). None of these behavioral changes were driven by changes in the locomotor activity as assessed by the OFT, and comparable behavioral response was observed between both groups (**Suppl Fig. 4**). Furthermore, we detected GI dysfunction characterized by increased gut motility time in DD-PD mice compared to the control group examined 8 weeks later (p < 0.05; **Fig. 4L**). At a cellular level, a marginal increase in the Iba1-ir signal (p = 0.0777) and significant increases in GFAP levels (p < 0.01) were detected in the ileum of DD-PD mice compared to controls (**Fig. 4M, 4N**). Together, these data and previous results confirm that the overexpression of α-Syn in raphe 5-HT neurons not only causes alterations in synaptic plasticity [23], but also induces changes in glia function that may contribute to the loss of integrity of the 5-HT system and its role in regulating emotional and autonomic functions, such as GI motility.

### Dysregulation of selected miRNA expression and inflammatory signaling pathways in the brain areas and ileum in DD-PD mice

To determine whether the changes in the miRNA-mRNA expression observed in human brain tissue were replicated in the DD-PD mouse model, we next examined the expression of the IBD-associated miRNA pattern and inflammatory markers in the mPFC, CPu, and ileum of mice 8 weeks after AAV-α-Syn infusion. Our analysis revealed a striking concordance between the human and animal data in both central and peripheral tissues. In the mPFC, a brain region that is highly conserved across different species and comparable to the human dlPFC, DD-PD mice exhibited significant downregulations of both miR-199a-5p (p < 0.01; **Fig. 5A**) and miR-219a-5p (p < 0.01; **Fig. 5B**) when compared with the control group. Conversely, miR-200a-3p was significantly upregulated (p < 0.05; **Fig. 5C**). A similar pattern was observed in the CPu, with significant reductions in miR-199a-5p (p < 0.05; **Fig. 5D**) and miR-219a-5p (p < 0.01; **Fig. 5E**), and a significant increase in miR-200a-3p (p < 0.05; **Fig. 5F**). To evaluate the gut-brain axis, we extended the analysis to include the ileum. Consistent dysregulation was also observed, characterized by a downward trend for miR-199a-5p (p = 0.0725; **Fig. 5G**), a significant decrease in miR-219a-5p (p < 0.05; **Fig. 5H**), and a substantial increase in miR-200a-3p (p < 0.01; **Fig. 5I**) in DD-PD mice compared to control group.

**Figure 5.**
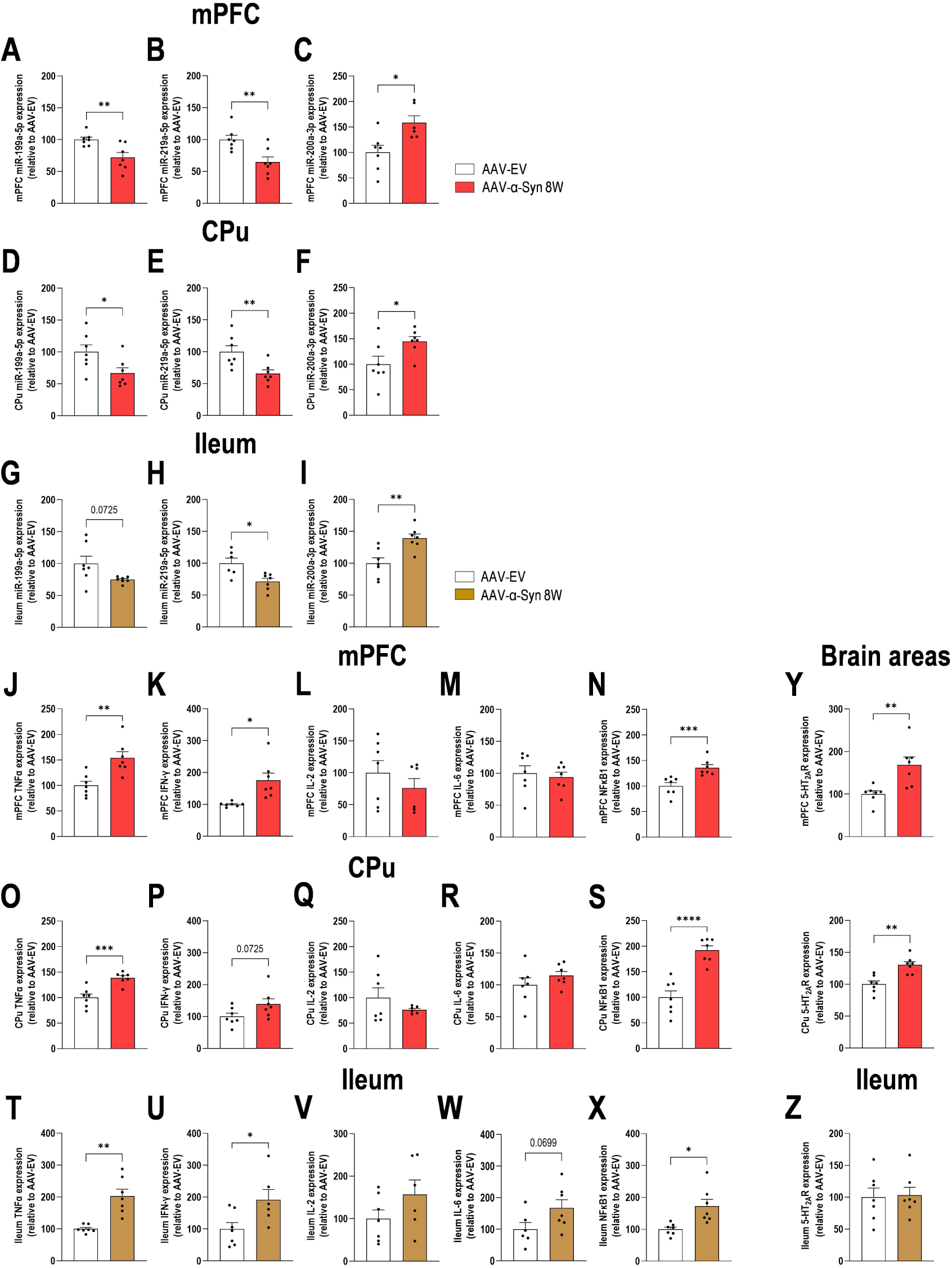
MiRNA and mRNA expression profile of inflammatory pathways in the mouse model of α-synucleinopathy with depressive phenotype. **A-C)** Expression levels of miR-199a-5p, miR-219a-5p, and miR-200a-3p in the medial prefrontal (mPFC) of mice overexpressing human α-Syn (AAV-α-Syn) in the serotonergic (5-HT) system compared to the control group (AAV-EV) at 8 weeks (W) post-infusion. **D-F)** Expression levels of these three miRNAs in the caudate-putamen (CPu) of the same AAV-α-Syn mice compared to AAV-EV mice. **G-I)** Expression levels of these three miRNAs in the ileum of the same AAV-α-Syn mice compared to AAV-EV mice. **J-N)** Expression levels of TNFα, IFN-γ, IL-2, IL-6, and NFκB1 mRNAs in the mPFC of the same AAV-α-Syn mice compared to AAV-EV mice. **O-X)** Expression levels of these inflammatory markers in CPu and ileum of the same AAV-α-Syn mice compared to AAV-EV mice. **Y-Z)** Expression levels of 5-HT_2A_R mRNA in mPFC, CPU, and ileum of AAV-α-Syn mice compared to AAV-EV mice. Data is plotted as mean ± SEM. Data points represent individual mice. * p < 0.05, ** p < 0.01, *** p < 0.001, **** p < 0.0001 versus controls (detailed statistical analysis in supplementary excel file).

Based on these findings, we investigated whether this miRNA profile was associated with a pro-inflammatory state in the same tissue samples. In the mPFC, DD-PD mice displayed significantly higher expression of TNFα (p < 0.01; **Fig. 5J**), IFN-γ (p < 0.05; **Fig. 5K**), and the transcription factor NFκB1 (p < 0.001; **Fig. 5N**). IL-2 and IL-6 expression did not change significantly (**Fig. 5L, 5M**), and values comparable to the control group were found. A consistent inflammatory profile was noted in the CPu, with significant increases in TNFα (p < 0.001; **Fig. 5O**) and NFκB1 (p < 0.0001; **Fig. 5S**). A trend towards increased IFN-γ expression was observed (p = 0.0725; **Fig. 5P**), while IL-2 and IL-6 levels remained unchanged (**Fig. 5Q, 5R**). Most importantly, this neuroinflammatory pattern was reflected in the ileum of the same DD-PD mice, which showed significant upregulation of TNFα (p < 0.01; **Fig. 5T**), IFN-γ (p < 0.05; **Fig. 5U**), and NFκB1 (p < 0.05; **Fig. 5X**). A marginal increase in IL-6 level was also observed (p = 0.0699; **Fig. 5W**), whereas IL-2 expression remained unaffected in the ileum of DD-PD mice (**Fig. 5V**) compared to controls.

In light of the relationship between the analyzed miRNAs and 5-HT signaling pathways, we also assessed 5-HT_2A_R mRNA expression in both the brain and ileum of the DD-PD mice. The analyses revealed increased 5-HT_2A_R expression in both the mPFC (p < 0.01) and CPu (p < 0.01), but not in in the ileum (**Fig. 5Y, 5Z**), of DD-PD mice compared to the control group. Overall, these results demonstrate that α-synucleinopathy originating in raphe 5-HT neurons is sufficient to trigger the parallel dysregulation of specific miRNAs and robust pro-inflammatory responses in various brain regions and the gut. This process recapitulates some of the molecular changes that we observed in post-mortem brains of patients with PD and DD.

### Changes in the expression of selected miRNAs and inflammatory marker mRNAs in the mPFC and CPu in a mouse model of corticosterone-induced depression

We then extended the analysis of IBD-associated miRNAs and their predicted mRNA targets, and included a depression-like mouse model based on chronic CORT consumption [43]. First, we confirmed the depressive phenotype by detecting longer immobility times in the FST (p < 0.05) and TST (p < 0.001) compared to the control group (**Fig. 6A, 6B**). As with brain samples from patients with DD, we detected downregulation of miR-199a-5p (p < 0.05 in the mPFC and CPu) and miR-219a-5p (p < 0.05 in the mPFC and p = 0.0558 in the CPu), as well as upregulation of miR-200a-3p (p < 0.05 in the mPFC and CPu), in mice exhibiting a depressive phenotype compared to the control group (**Fig. 6C-H**).

**Figure 6.**
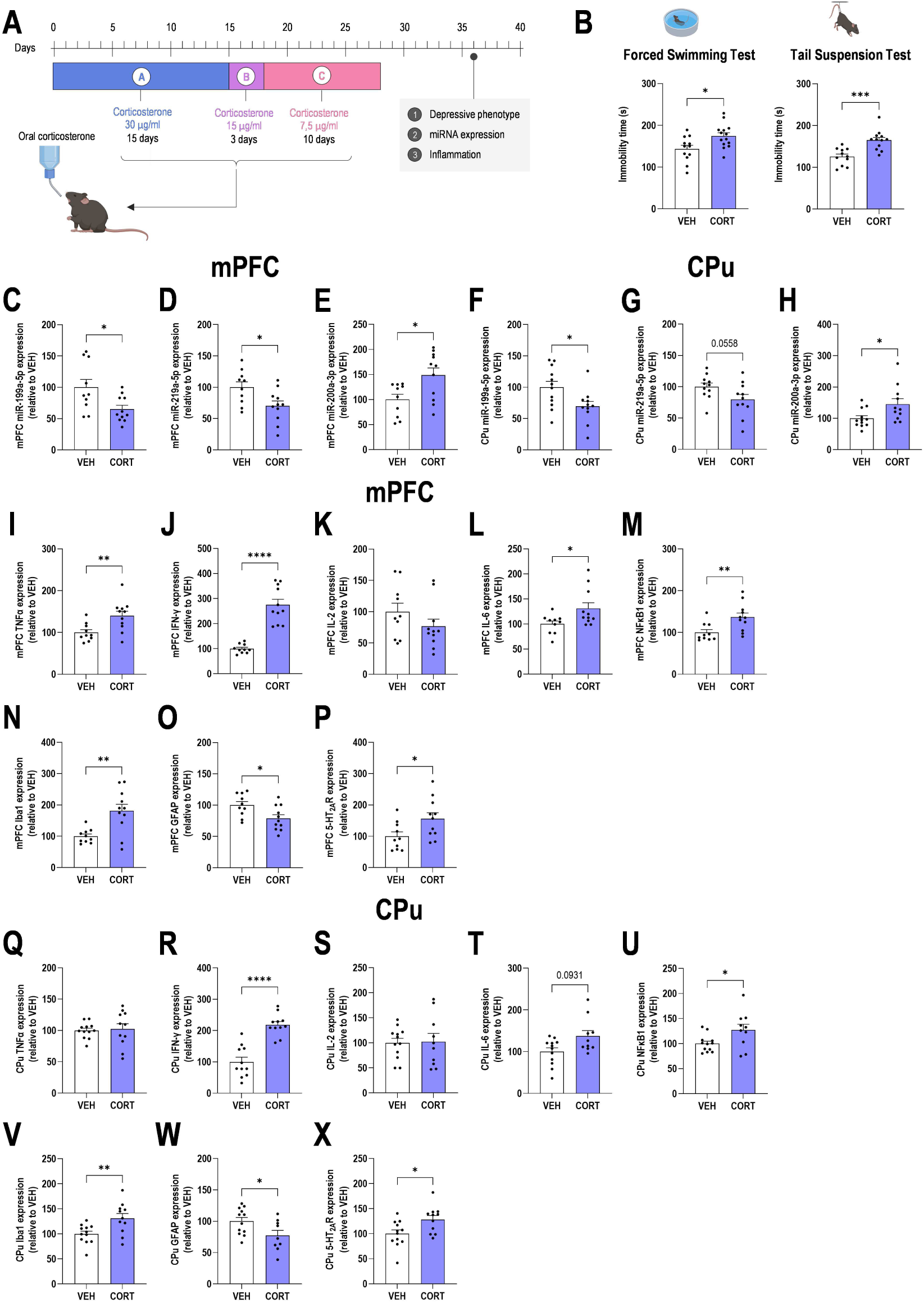
MiRNA and mRNA expression profile in a depressive-like mouse model subjected to corticosterone (CORT)-induced stress. **A)** Experimental design. Mice were chronically CORT exposure in the drinking water for 28 days and examined on day 36 [42]. **B)** Mice exposed to CORT displayed depressive-like behaviors, including increased immobility time in the forced swimming and tail suspension tests. **C-H)** Expression levels of miR-199a-5p, miR-219a-5p, and miR-200a-3p in the medial prefrontal (mPFC) and caudate putamen (CPu) of CORT-treated mice compared to the control vehicle (VEH)-treated mice. **I-J)** Expression levels of TNFα, IFN-γ, IL-2, IL-6, and NFκB1 mRNAs in the mPFC of the same CORT-treated mice compared to the VEH-treated mice. **N-P)** Expression levels of Iba1, GFAP, and 5-HT_2A_R mRNAs in the mPFC of the same CORT-treated mice compared to the VEH-treated mice. **Q-U)** Expression levels of TNFα, IFN-γ, IL-2, IL-6, and NFκB1 mRNAs in the CPU of CORT-treated mice and VEH-treated mice. **V-X)** Expression levels of Iba1, GFAP, and 5-HT_2A_R mRNAs in the CPu of CORT-treated mice and VEH-treated mice. Data is plotted as mean ± SEM. Data points represent individual mice. * p < 0.05, ** p < 0.01, *** p < 0.001, **** p < 0.0001 versus controls (detailed statistical analysis in supplementary excel file).

Analysis of the expression of inflammatory target mRNAs revealed significant increases in TNFα (p < 0.01, **Fig. 6I**), IFN-γ (p < 0.0001, **Fig. 6J**), IL-6 (p < 0.05, **Fig. 6L**), and NFκB1 (p < 0.01, **Fig. 6M**) in the mPFC of CORT-treated mice compared to the controls. At the same time, we observed a significant increase in IFN-γ expression (p < 0.0001, **Fig. 6R**) and in NFκB1 expression (p < 0.05, **Fig. 6U**), as well as a marginal increase in IL-6 levels (p = 0.0931, **Fig. 6T**) in the CPu of CORT-treated mice compared to the control group.

Finally, mice exhibiting a depressive phenotype showed a significant increase in Iba1 mRNA expression in the mPFC (p < 0.01) and CPu (p < 0.01), as well as a decrease in GFAP mRNA levels in the mPFC (p < 0.05) and CPu (p < 0.05), when compared to the control group (**Fig. 6N, 6V, 6O, 6W**). In addition, a significant increase was also detected in the 5-HT_2A_R mRNA expression in both mPFC and CPu (p < 0.05) of depressive-like mice (**Fig. 6P, 6X**). Overall, these results highlight that the IBD-associated miRNAs, namely miR-199a-5p, miR-219a-5p, and miR-200a-3p, as well as target mRNAs of NFκB1-downstream inflammatory pathways and 5-HT signaling, are significantly affected in PD, DD, and its comorbidity.

## Discussion

There is growing evidence linking PD and DD to intestinal inflammation, also known as leaky gut syndrome, through a shared pathophysiology involving abnormal miRNA expression [40, 41]. Patients with IBD are at much higher risk of developing PD and/or DD, and *vice versa*, than the general population [25–29]. In this study, we identified a conserved molecular miRNA pattern that links the pathophysiology of PD, DD and GI dysfunction. Our findings demonstrate that a specific triad of miRNAs —downregulated miR-199a-5p and miR-219a-5p, and upregulated miR-200a-3p— is a common feature in the post-mortem brains of PD (both early B2-3 and late B5-6 stages) and DD patients, as well as being present in patients with IBD [40]. This profile of dysregulated miRNAs is closely associated with the direct activation of the NFκB1 mRNA target and a pro-inflammatory state characterized by increased expression of mainly TNFα and IFN-γ cytokines, as well as Iba1-dependent microglia activation in PD and DD human brain samples. Importantly, we have shown that this molecular cascade occurs in two mouse models associated with (i) CORT-induced stress and (ii) α-synucleinopathy affecting the 5-HT system, inducing a depressive-like phenotype. In the latter, changes in miRNA-mRNA expression occur in both the brain areas and the ileum, providing strong evidence that the gut-brain axis is disrupted in these concurrent disorders. It should be noted that GO and KEGG analysis based on the three selected miRNAs showed an enrichment of the 5-HT pathway, and we confirmed increases in 5-HT_2A_R mRNA expression in both brain samples from PD and DD patients and in both mouse models, adding further evidence of a common pattern of dysregulated miRNAs that converge in both pathologies.

Identifying these three specific miRNAs (miR-199a-5p, miR-219a-5p, and miR-200a-3p), which have been previously described in IBD [52–54], could provide a molecular link between the high clinical prevalence of PD, DD, and GI dysbiosis in patients. Our bioinformatic analysis revealed that the predicted targets of these miRNAs converge on enriched signaling pathways, including PI3K-Akt (hsa04151), MAPK (hsa04010), Wnt (hsa04310), TNF (hsa04468), and neurotrophin (hsa04722), among others. These pathways are master regulators of cellular processes frequently disrupted in both PD and DD, such as synaptic plasticity, cellular stress responses, and, most notably, neuroinflammation and immune responses. In fact, the IBD signaling pathway (hsa05321) itself appears among the top 10 most enriched pathways in the KEEG analysis. Among the numerous miRNAs dysregulated in IBD, miR-199a-5p, miR-219a-5p, and miR-200a-3p, consistently emerge as miRNAs whose expression is significantly altered in the presence of active inflammation [40, 55]. Several reports indicate that miR-219a-5p is significantly downregulated in patients with IBD, as we have reported here in the dlPFC and CAU of patients with PD and DD. It has been suggested that the inflammatory microenvironment, including IL-6, IL-12, IL-23, and TNFα, directly inhibits the expression of miR-219a-5p in CD4^+^ T cells, which are key drivers of the adaptive immune response in IBD [56]. Furthermore, it has been reported that miR-219a-5p plays a broader role in maintaining overall intestinal homeostasis and barrier function, and its loss contributes to a number of intestinal disorders [57]. Our findings are consistent with these data, as we systematically observed increased expression of several cytokines (e.g., TNFα, IFN-γ, IL-2, and IL-6) in post-mortem samples from patients with PD and DD, as well as in mouse models with PD and DD phenotypes, depending on the brain area analyzed. Although data on miR-219a-5p expression in PD patients are scarce, it has been reported that plasma levels of a closely related isoform, miR-219a-2-3p, are decreased in a PD rat model compared to controls [58]. Similarly, there is very little data on the role of miR-219a-5p in DD. Instead, a study using postmortem dlPFC samples from subjects with DD identified differential regulation of miR-219a-2-3p in the synaptic fraction compared to healthy subjects [59]. Although both miRNAs may originate from the same precursor gene (hsa-mir-219a-2), miR-219a-5p and miR-219a-2-3p are distinct mature molecules and, consequently, may regulate completely different sets of target genes. Taken together, our findings emphasize the importance of miR-219a-5p in the pathophysiology of PD and DD. In addition to its well-documented roles in regulating myelination [60] and modulating glutamatergic signaling via the NMDA receptor [61], our data suggest that it also plays a significant part in the inflammatory response.

In contrast, the situation with miR-199a-5p and miR-200a-3p in IBD is more complex. This is evident from their opposing expression levels, which depend largely on the type of tissue analyzed (e.g. ileum, peripheral blood, or fecal matter) [47, 62–65]. Nevertheless, the effects of miR-199a-5p on IBD pathogenesis occur through at least two interconnected mechanisms. The first involves altering the intestinal epithelial barrier by regulating the Rho-associated protein kinase (ROCK) signaling pathway, which plays a key role in modulating the actin cytoskeleton, cell adhesion, and tight junction integrity [66, 67]. The second mechanism involves directly amplification of inflammatory signaling pathways though the NFκB1 pathway [68]. Indeed, *in vitro* assays using luciferase reporter demonstrated that miR-199a-5p binds to the 3’-UTR of NFκB1 mRNA and therefore regulates NFκB1 function (miRTarBase, #MIRT438174). Although we did not perform reporter assays, we validated NFκB1 mRNA expression and observed consistent increases in its expression in the dlPFC and CAU of PD and DD patients, as well as in both mouse models, suggesting direct regulation of its transcription mediated by miR-199a-5p. It should be noted that the activation of the NFκB1 pathway increases the production of pro-inflammatory cytokines, including TNFα, IFN-γ, among others, in IBD mouse models [68]. Similarly, we also detected increases in the expression of these cytokines in human brain samples with PD or DD and in brain and ileum samples in the PD mouse model with depressive-like phenotype. Therefore, this orchestrated change in cytokine balance creates a potent and self-sustained pro-inflammatory environment that may link gut-brain axis disorders, including PD and DD.

In the context of PD, previous data also support a decrease in miR-199a-5p expression in DA neurons generated from induced pluripotent stem cells (iPSCs) from sporadic and LRRK2 mutant patients [69], providing strong evidence for the involvement of miR-199a-5p in the pathophysiology of PD. In fact, it has been suggested that miR-199a-5p function is related to its potent regulation of the apoptosis mechanism. Previous studies demonstrated that miR-199a-5p directly targets the mRNA of the X-linked inhibitor of apoptosis protein (XIAP) [70]. The apoptosis of DA neurons is a central event in the pathology of PD, making both the regulation of miR-199a-5p/XIAP and miR-199a-5p/NFκB1 described here highly relevant to the mechanism of the disease. Furthermore, contrary to the reduction in miR-199a-5p levels in the dlPFC and CAU of patients with DD, a previous study has shown an increase in miR-199a-5p expression in the serum and cerebrospinal fluid of patients with DD compared to non-depressed controls [71]. However, this study has the limitation of having a small sample size of seven subjects. The same authors also reported increased miR-199a-5p expression in the hippocampus of male mice that were subjected to chronic unpredictable mild stress [71], in contrast to decreased miR-199a-5p expression in the mPFC and CPu in female mice chronically exposed to CORT (this study). Although further studies are needed to unravel the sex- and tissue-dependent role of miR-199a-5p in the pathophysiology of DD, experimental validation using luciferase reporter assays has confirmed that miR-199a-5p also binds directly to the 3’UTR of WNT2 mRNA (miRTarBase - #MIRT438595), resulting in an altered downstream cascade. WNT signaling normally leads to the activation and phosphorylation of the cAMP-response element binding protein (CREB), a transcription factor that plays an essential role in neuronal plasticity and survival. Therefore, changes in phosphorylated CREB levels (p-CREB) are to be expected when the regulation of WNT2 by miR-199a-5p is disrupted. This is critical, as p-CREB drives the transcription of several key neurotrophic genes, such as brain-derived neurotrophic factor (BDNF), which is directly related to the DD pathophysiology [72].

As for miR-200a-3p, although some studies showed a clear downregulation of miR-200a-3p and other members of the miR-200 family (e.g., miR-200b and miR-200c) in the inflamed colonic tissue of IBD patients [62] and in the plasma of DD patients [73]; most reports indicated a marked miR-200a-3p upregulation in the blood/serum of patients with both inflammatory ulcerative colitis and Crohn’s disease compared to healthy individuals [64]. Similarly, upregulation of miR-200a-3p expression has been observed across different experimental paradigms, from *in vitro* cell models to human PD patient samples and in differentiated PC12 cells treated with the neurotoxin 1-methyl-4-phenylpyridinium (MPTP) [74]. In addition, evidence points to an increase in miR-200a-3p in the dlPFC and serum of patients diagnosed with DD compared to controls [75, 76]. Our data are consistent with these observations and add solid information on miR-200a-3p upregulation in different brain areas of patients with PD and DD, as well as in both mouse models of DD-PD comorbidity and CORT-induced stress. It has been reported that overexpression of miR-200a-3p promotes inflammation by inducing the activation of the NLR family pyrin domain containing 3 (NLRP3) inflammasome [77]. Similarly, several studies have established that sirtuin 1 (SIRT1) is a direct target of miR-200a-3p [78, 79]. SIRT1 is a highly conserved NAD^+^-dependent protein deacetylase that functions as a master regulator of cellular metabolism, stress resistance, and longevity [80]. Within the brain, SIRT1 has potent and multifaceted neuroprotective functions that are directly relevant to the pathology of PD [81], and more recently linked to DD [82]. Therefore, changes in miR-200a-3p/SIRT1 regulation may affect mitochondrial health and biogenesis, the clearance of misfolded proteins (such as α-Syn) and neuroinflammatory pathways in PD/DD.

Additionally, some studies have shown that miR-200a-3p regulates genes within the 5-HT pathway, as its direct binding to *SLC6A4* (SERT) has been reported. Indeed, one factor that exacerbates visceral hyperalgesia and hypersensitivity is the suppression of SERT mRNA by miR-200a-3p [40]. Although bioinformatic analysis of the three miRNAs examined in this study predicts potential binding sites for both miR-200a-3p and miR-199a-5p in *HTR2A* (5-HT_2A_R), this interaction has not yet been confirmed by standard luciferase reporter assays. Nevertheless, we validated 5-HT_2A_R as a target and found elevated levels of 5-HT_2A_R mRNA in the same brain regions affected in PD and DD (as well as in mouse models of DD-PD comorbidity and CORT-induced stress), where levels of miR-200a-3p and miR-199a-5p were altered. While we cannot establish a direct causal relationship between the described miRNAs and their target mRNAs, our results suggest that the alteration of serotonergic signaling through 5-HT_2A_R is present in both PD and DD. In fact, the density of 5-HT_2C_R and 5-HT_2A_R in the SN pars reticulate and CAU appears to increase in PD patients [83, 84]. Similarly, several studies have also demonstrated alterations in 5-HT_2A_R density in the brain of depressed subjects [85, 86].

A detailed analysis of the glial markers Iba1 and GFAP reveals nuanced differences in the neuroinflammatory response, suggesting that while inflammation is a common feature in PD and DD, its cellular characteristic is context-dependent. Iba1 expression, a marker of microglial activation, was significantly increased in the brains of late-stage PD patients, DD patients, DD-PD mice (in the DR, mPFC, CPu, as well as in the ileum), and in CORT-exposed mice (in the mPFC and CPu). This establishes microglial activation as a consistent and central feature of the neuroinflammation observed in all the conditions studied. Microglia appears to play a role in responding to both proteopathic (such as α-Syn toxic seeding in PD) and stress-related insults [87–90]. Furthermore, the temporal data from the DD-PD mouse model, show a stronger microglial response at 4 weeks than at 8 weeks, leading to neuroinflammatory cascade that may contribute to the dysregulation of the miR-199a-5p, miR-219a-5p, miR-200a-3p, and others [91]. Recent data have shown that microglial activation acts as an initial barrier against aggregated α-Syn. However, when the phagocytic/degradative function of microglia is overwhelmed, their ability to provide neuroprotection diminishes, leading to the spread of α-Syn between cells and exacerbating the inflammatory process [92]. Unlike the similar microglial response, the expression of GFAP, a marker of reactive astrocytes, differs. While GFAP expression is upregulated in the dlPFC and CAU in late-stage PD patients and in the DD-PD mouse model (in the DR, mPFC, and ileum); it is downregulated in the dlPFC of DD patients and, in the mPFC and CPu of the CORT mouse model. This is a critical finding that distinguishes the nature of neuroinflammation in the two disorders. Our data are consistent with previous findings showing that α-Syn pathology induces reactive and hypertrophic astrocytes with increased GFAP signaling. This is likely to be a response to failures in the autophagy and lysosome pathways, which have been shown to impair α-Syn clearance in astrocytes [93–95]. On the other hand, multiple brain regions from individuals with major DD have lower expression levels of astrocyte markers (e.g., GFAP, glutamine synthase - GS, among others) and lower densities of astrocytes labeled for these marker [96–98]. This astrocytic atrophy may suggest a loss of crucial neuroprotective functions and synaptic plasticity, which are increasingly linked to the depression pathophysiology [99]. Therefore, systematic findings obtained from human brain samples of patients with PD and DD, as well as from murine models of both disorders, provide a new biological basis for distinguishing neuroinflammation in DD-PD comorbidity from DD alone. This suggests that, while microgliosis is a common factor, the degree of astrocytic involvement is a key differentiating factor.

Finally, the mouse model with α-synucleinopathy specific to the 5-HT system validates the relationship between PD and the presence of non-motor symptoms. The mice developed a robust depressive phenotype [22, 23] and GI dysmotility without motor deficits, reflecting the early non-motor symptoms of PD. This behavioral phenotype is supported by significant central neuroinflammation, evidenced by pronounced microglia and astrogliosis in key brain regions such as the DR, mPFC and CPu, suggesting that toxic α-Syn accumulation originated in the 5-HT system is sufficient to trigger a generalized inflammatory response. A relevant aspect was to demonstrate that the effects associated with α-Syn extend beyond the brain. The replication of the same profile of miRNAs and inflammatory cytokines in the ileum of the DD-PD mice provides direct evidence of gut-brain axis communication. The presence of elevated levels of TNFα, IFN-γ, and NFκB1 in both the gut and brain of the same animal suggests a shared, self-perpetuating inflammatory loop in which peripheral inflammation may drive central pathology, and vice versa. Recent studies, using a mouse model that overexpresses the mutant A53T α-Syn in DA neurons of the SNc, demonstrated the presence of aggregated α-Syn in the ileum within CD11c^+^ microglia-like cells, providing a mechanism for α-Syn trafficking between the brain and gut [44].

In conclusion, this study identifies a conserved inflammatory-miRNA axis comprising miR-199a-5p, miR-219a-5p and miR-200a-3p that is dysregulated in PD, DD and associated intestinal pathology. The upregulation of proinflammatory miR-200a-3p, together with the downregulation of miR-199a-5p and miR-219a-5p (which are often involved in anti-inflammatory and neuroprotective functions), likely creates a sustained and unresolved inflammatory environment. This environment, rich in cytotoxic mediators such as TNFα and IFN-γ, can impair synaptic function, alter neurotransmitter homeostasis, and ultimately contribute to the neurodegenerative process and the manifestation of depressive symptoms. Constant activation of NFκB1, a master regulator of the inflammatory response, in all affected tissues underscores its central role in coordinating this common pathological cascade (**Table 2**). Therefore, this pattern represents a critical point of molecular convergence, driving a shared neuroinflammatory process spanning the gut-brain axis. Targeting this triad of miRNAs could represent a novel, transformative therapeutic strategy with the potential to alleviate the motor, psychiatric and gastrointestinal symptoms of these devastating disorders simultaneously.

**Table 2.** Summary of key molecular changes across human cohorts and preclinical mouse models.

| Marker | PD human<br>(dlPFC/CAU) | DD human<br>(dlPFC/CAU) | DD-PD mouse<br>(mPFC/CPu) | DD-PD mouse<br>(Ileum) | CORT mouse<br>(mPFC/CPu) |
| --- | --- | --- | --- | --- | --- |
| miR-199a-5p | ↓ | ↓ | ↓ | ↓ | ↓ |
| miR-219a-5p | ↓ | ↓ | ↓ | ↓ | ↓ |
| miR-200a-3p | ↑ | ↑ | ↑ | ↑ | ↑ |
| NFκB1 | ↑ | ↑ | ↑ | ↑ | ↑ |
| TNFα | ↑ | ↑ | ↑ | ↑ | ↑ (mPFC) |
| INF-γ | ↑ | ↑ (CAU) | ↑ | ↑ | ↑ |
| Iba1 | ↑ | ↑ | ↑ | ↑ | ↑ |
| GFAP | ↑ | ↓ | ↑ | ↑ | ↓ |
| 5-HT<sub>2A</sub>R | ↑ | ↑ | ↑ | - | ↑ |

Note: ↑ indicates upregulation; ↓ indicates downregulation; - indicated unchanged

### Limitations and future directions

Although our findings provide a robust and consistent picture, certain limitations must be acknowledged. Firstly, human data derived from post-mortem tissue are inherently correlational and cannot establish causality. Furthermore, while DD-PD mouse model is highly relevant for studying non-motor symptoms related to the 5-HT system, it does not fully encompass the progressive DA degeneration characteristic of advanced PD. The CORT model used female mice, whereas the DD-PD model used males. Although data from human DD brains, but not in PD cohort, were analyzed by sex, further studies are needed to fully explore sex-dependent effects in the models. Future studies should validate this miRNA pattern as a potential biomarker in patients’ biological fluids (e.g. plasma or cerebrospinal fluid), as this could be a powerful tool for early diagnosis and patient stratification. Additionally, future research should investigate the specific roles of these miRNAs in microglia, astrocytes and neurons to determine their exact contribution to the inflammatory cascade.

## Author contributions

**LMR** contributed to the conceptualization and experimental design of the study, performed human/mouse miRNA and mRNA analyses: qPCR, GO, KEGG pathways, and statistical analyses, and co-wrote and edited the manuscript. **JJE** contributed to the experimental design, and performed confocal microscopy, IMARIS 3D images acquisition, and statistical analyses. **CYC and USS** participated in the mouse models, behavioral assessment, and preparation of brain and ileum tissue samples. **VP** was involved with tissue preparation and immunohistochemistry. **LFC and JJM** participated in the collection of human tissue data. **AB** contributed to the conceptualization and experimental design of the study, co-wrote and edited the manuscript, supervised the entire study and provided the necessary resources. All authors reviewed the manuscript. All authors read and approved the final manuscript.

## Ethical statement

Postmortem human brain samples

The study was approved by the Ethics Committee of Spanish National Research Council (Project Ref. PID2019-105136RB-I00). The dorsolateral prefrontal cortex (dlPFC) and caudate nucleus (CAU) tissue samples from controls and PD patients were obtained from the Hospital Clinic-IDIBAPS biobank, in accordance with Spanish legislation and with the approval of the local ethics committees. Pathological cases were categorized as stages 1–6 of LB disease pathology, according to Braak staining (Braak et al., 2003), excluding tauopathies, vascular disease, and metabolic syndrome. Cases in the control group had not neurological, psychiatric, or metabolic disorders and no neuropathological abnormalities, except for sporadic Alzheimer’s disease. A total of 10 control and 20 cases with PD-related pathology were included in the present study. All PD cases were treated for motor symptoms. See **Table 1** for subject details.

In addition, dlPFC and CAU tissue samples were obtained at autopsy from non-psychiatric controls and DD patients at the Basque Institute of Legal Medicine, Bilbao, Spain, in accordance with the Spanish guidelines of research and ethics committees for postmortem brain studies. This study used dlPFC and CAU from a total of 28 controls and 28 subjects with DD. Of the 28 subjects with MDD, 23 died by suicide. Cohort characteristics are shown in **Table 1**.

