## Supplemental Figures for "A molecular convergence in the triad of Parkinson’s disease, depressive disorder and gut health is revealed by the inflammation-miRNA axis"

### Supplementary Figure 1

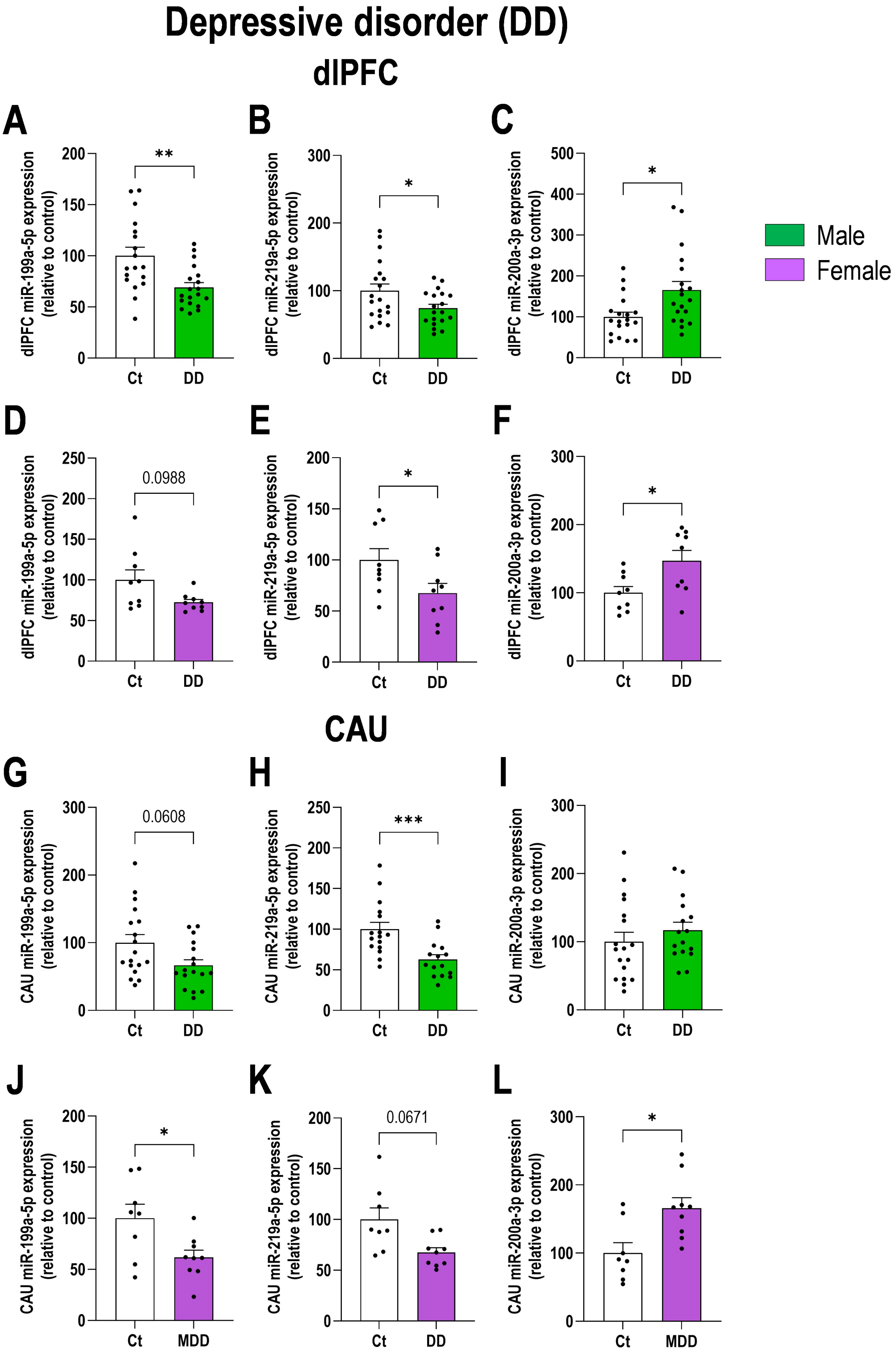


**Supplementary Figure 1.** Sex-dependent analysis of dysregulated miRNAs in postmortem brains samples of patients with depressive disorder (DD). **A-C)** Expression levels of miR-199a-5p, miR-219a-5p, and miR-200a-3p in the dorsolateral prefrontal cortex (dlPFC) of male patients with DD compared to non-DD males in the control group (Ct). **D-F)** Expression levels of these three miRNAs in the dlPFC of female patients with DD compared to non-DD female in the Ct group. **G-H)** Expression levels of miR-199a-5p, miR-219a-5p, and miR-200a-3p in the caudate nucleus (CAU) of male patients with DD compared to non-DD males in the Ct group (Ct). **J-L)** Expression levels of these three miRNAs in the CAU of female patients with DD compared to non-DD female in the Ct group. Data is plotted as mean ± SEM. Data points represent individual human samples. * $p < 0.05$, ** $p < 0.01$, *** $p < 0.001$ versus respective controls (detailed statistical analysis in supplementary excel file).

### Supplementary Figure 2

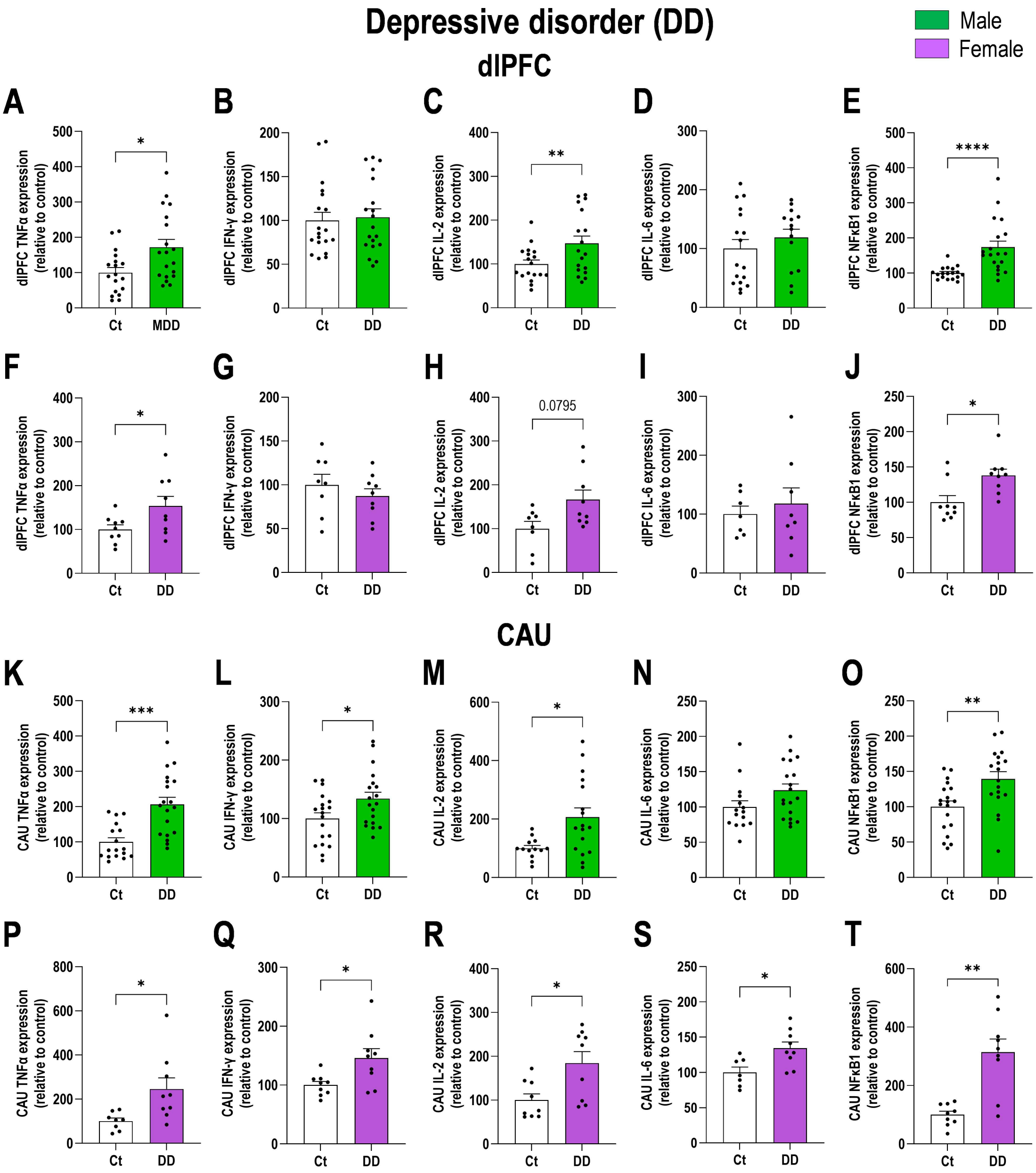


**Supplementary Figure 2.** Sex-dependent analysis of inflammatory mRNAs markers in postmortem brains samples of patients with depressive disorder (DD). **A-E)** Expression levels of TNFα, IFN-γ, IL-2, IL-6, and NFκB1 mRNAs in the dorsolateral prefrontal cortex (dlPFC) of male patients with DD compared to non-DD males in the control group (Ct). **F-J)** Expression levels of the same inflammatory marker mRNAs in the dlPFC of female patients with DD compared to non-DD female in the Ct group. **K-O)** Expression levels of TNFα, IFN-γ, IL-2, IL-6, and NFκB1 mRNAs in the caudate nucleus (CAU) of male patients with DD compared to non-DD males in the Ct group (Ct). **J-L)** Expression levels of the same inflammatory marker mRNAs in the CAU of female patients with DD compared to non-DD female in the Ct group. Data is plotted as mean ± SEM. Data points represent individual human samples. * $p < 0.05$, ** $p < 0.01$, *** $p < 0.001$, **** $p < 0.0001$ versus respective controls (detailed statistical analysis in supplementary excel file).

**Supplementary Figure 3**

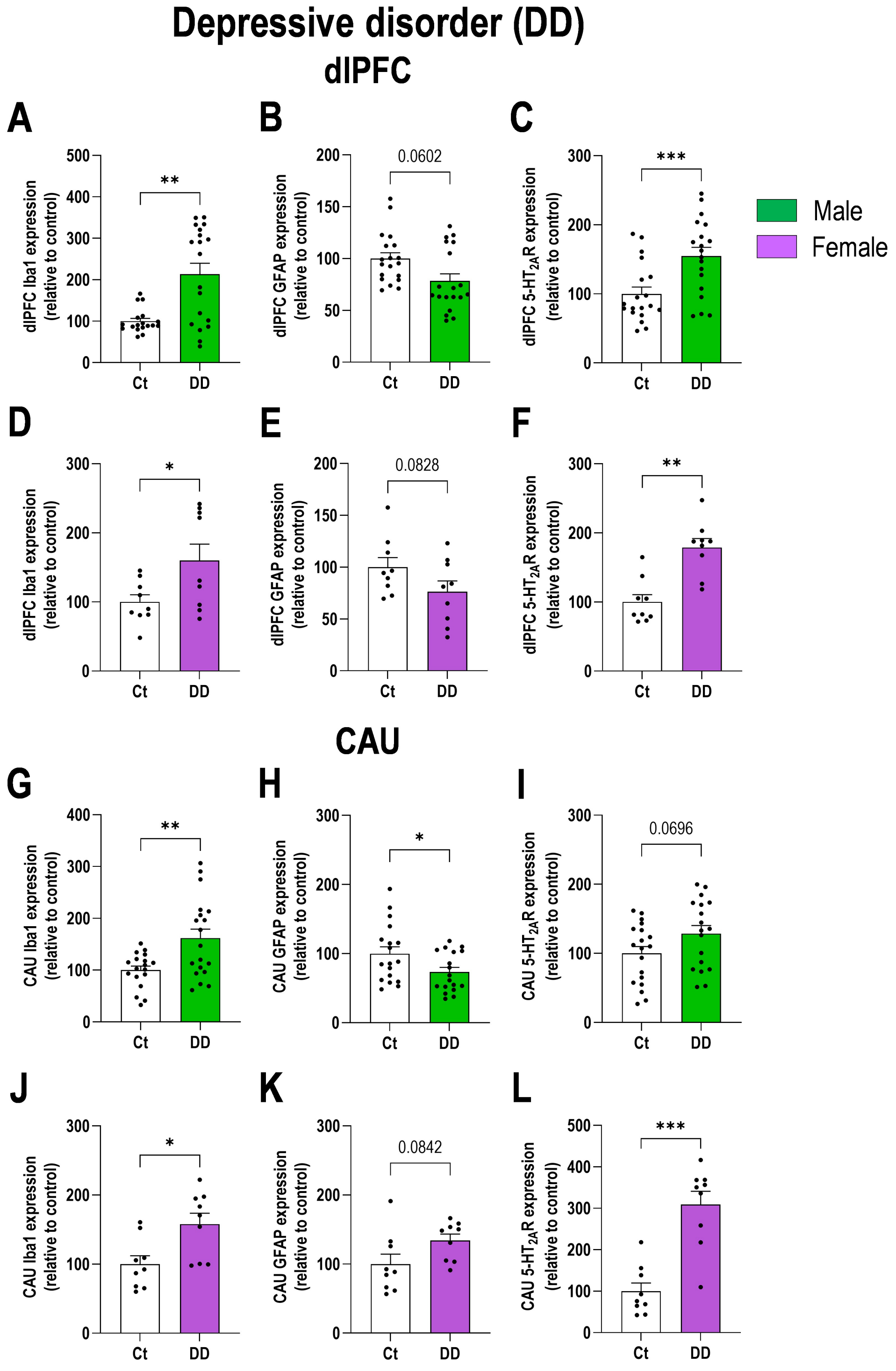


**Supplementary Figure 3.** Sex-dependent analysis of glial and serotonin (5-HT) signaling markers in postmortem brains samples of patients with depressive disorder (DD). **A-C)** Expression levels of Iba1, GFAP, and 5-$HT_{2A}$R mRNAs in the dorsolateral prefrontal cortex (dlPFC) of patients with DD compared to non-DD males in the control group (Ct). **D-F)** Expression levels of the same mRNAs in the dlPFC of female patients with DD compared to non-DD female in the Ct group. **G-I)** Expression levels of Iba1, GFAP, and 5-$HT_{2A}$R mRNAs in the caudate nucleus (CAU) of male patients with DD compared to non-DD males in the Ct group (Ct). **J-L)** Expression levels of the same mRNAs in the CAU of female patients with DD compared to non-DD female in the Ct group. Data is plotted as mean ± SEM. Data points represent individual human samples. * $p < 0.05$, ** $p < 0.01$, *** $p < 0.001$ versus respective controls (detailed statistical analysis in supplementary excel file).

**Supplementary Figure 4**

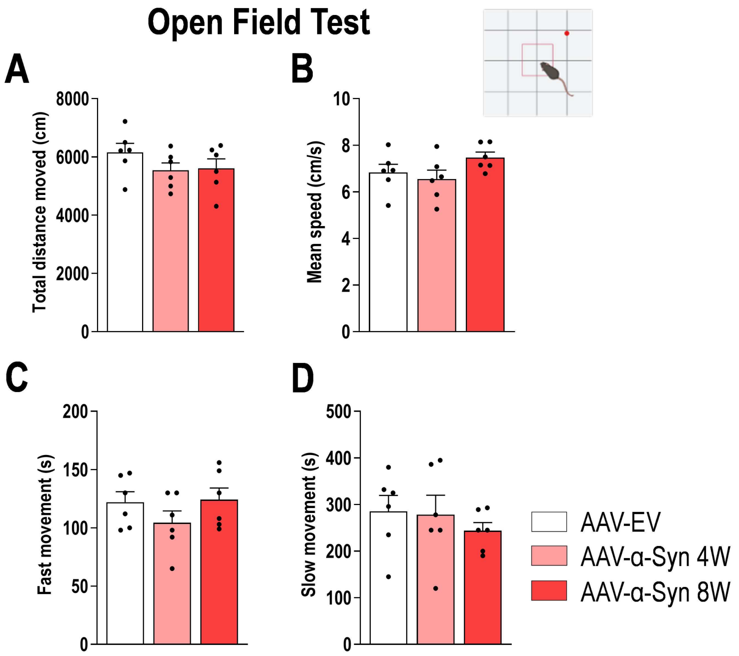


**Supplementary Figure 4.** Locomotor activity in the mouse model of α-synucleinopathy with depressive phenotype. **A-B)** Mice with human α-Syn accumulation in the serotonin (5-HT) system displayed comparable performance in the open field compared with control mice (AAV-EV). Bar graph showing quantification of: total distance traveled (A), average speed (B), fast movement (C), and slow movement (D). Data is plotted as mean ± SEM. Data points represent individual mice (detailed statistical analysis in supplementary excel file).
